# CDH3-AS1 antisense RNA enhances P-cadherin translation and acts as a tumor suppressor in melanoma

**DOI:** 10.1101/2024.12.26.630428

**Authors:** Manon Chadourne, Crystal Griffith, Xiaonan Xu, Emily Brennan, Olga Vera, Nicol Mecozzi, Kaizhen Wang, Alex M. Jaeger, Florian A. Karreth

**Author notes:** Present address: Biomarkers and Experimental Therapeutics in Cancer, IdiPAZ, and Cancer Epigenetics Laboratory, INGEMM, La Paz University Hospital, Madrid 28046, Spain (Olga Vera). **Conflicts:** The authors declare no financial conflicts of interest.

## Abstract

Thousands of regulatory noncoding RNAs (ncRNAs) have been annotated; however, their functions in gene regulation and contributions to cancer formation remain poorly understood. To gain a better understanding of the influence of ncRNAs on gene regulation during melanoma progression, we mapped the landscape of ncRNAs in melanocytes and melanoma cells. Nearly half of deregulated genes in melanoma are ncRNAs, with antisense RNAs (asRNAs) comprising a large portion of deregulated ncRNAs. *CDH3-AS1*, the most significantly downregulated asRNA, overlaps the *CDH3* gene, which encodes P-cadherin, a transmembrane glycoprotein involved in cell adhesion that was also reduced in melanoma. Overexpression of *CDH3-AS1* increased cell aggregation and reduced xenograft tumor growth, mimicking the tumor-suppressive effects of *CDH3*. *CDH3-AS1* interacted with *CDH3* mRNA and enhanced P-cadherin protein levels. Interestingly, secondary structures at the *CDH3* 5’ end regulated P-cadherin translation, and ribosome profiling revealed that *CDH3-AS1* promotes ribosome occupancy at the *CDH3* mRNA. Notably, ribosome occupancy was generally increased in mRNAs having cognate asRNA that are complementary to the 5’UTR. Taken together, this study revealed the *CDH3-AS1*-mediated enhancement of P-cadherin translation, underscoring the broader potential of asRNAs as regulators of protein-coding genes and their role in diseases like melanoma.

## INTRODUCTION

Melanoma is a type of cancer originating from melanocytes and although metastatic melanoma accounts for only 1% of all skin cancer cases, it causes the majority of skin cancer-related deaths (1). Genetic and genomic alterations are well-established drivers of melanoma initiation and progression, with oncogenic mutations in BRAF, NRAS, or NF1 being the initiating events, leading to hyperactivation of the MAPK pathway (2). Targeted therapies for melanoma primarily inhibit the MAPK pathway; however, the emergence of resistance (3–5) underscores the need for a deeper understanding of melanoma development to identify new potential therapeutic targets.

Initially considered transcriptional noise, non-coding RNAs (ncRNAs) are now recognized as major regulators of gene expression, playing critical roles in human development and diseases, including cancer (6–9). ncRNAs are classified based on their length into small ncRNAs (shorter than 200 nucleotides) and long ncRNAs (lncRNAs, longer than 200 nucleotides). lncRNAs constitute the largest portion of the mammalian non-coding transcriptome (7). One subclass of lncRNAs are antisense ncRNAs (asRNAs, also termed Natural Antisense Transcripts - NATs), which are transcribed from the opposite strand of cognate sense transcripts, regardless of whether the sense transcript encodes a protein. Previous studies showed that approximately a quarter of human protein-coding genes have a corresponding asRNA (10, 11), suggesting the potential for a significant proportion of protein coding genes to be regulated by asRNAs. asRNAs can influence nearly all aspects of gene expression, from transcription and translation to RNA degradation, either through the act of asRNA transcription or through various mechanisms involving the transcribed asRNA molecule (12). Multiple asRNAs have been shown to play roles in cancer development and progression (13, 14). While some lncRNAs have been shown to influence the development of melanoma (15–18), the roles of asRNAs in this malignancy are poorly understood.

In this study, we profiled the long noncoding RNA landscape in melanoma, identifying numerous deregulated lncRNAs including many asRNAs. We characterized the regulation and function of the most downregulated asRNA in human melanoma cells, *CDH3-AS1* (RP11-615I2.2). *CDH3- AS1* overlaps the protein-coding gene *CDH3*, which encodes the cell-cell adhesion protein P-cadherin, a putative tumor suppressor that is also significantly downregulated in melanoma cells. We demonstrated that both genes are negatively regulated by the MAPK pathway and elicit tumor suppressor effects. Additionally, we demonstrated that *CDH3-AS1* regulates P-cadherin translation, primarily in cis, through its interaction with *CDH3* mRNA. Secondary structures were identified at the 5’ end of *CDH3*, corresponding to the region overlapping with *CDH3-AS1*. Ribosome profiling revealed that ribosome occupancy on *CDH3* is influenced by *CDH3-AS1*. Furthermore, we observed a general increase in ribosome occupancy in mRNAs having complementary asRNAs. This study uncovers a potential mechanism by which asRNAs enhance translation efficiency, highlighting the regulatory role of asRNA in cancer formation.

## MATERIALS AND METHODS

### Cell Culture and Treatments

Human melanocyte cell lines (Hermes1, Hermes2, Hermes3A, and Hermes4B) were obtained from St. George’s University of London (https://www.sgul.ac.uk/about/our-institutes/neuroscience-and-cell-biology-research-institute/genomics-cell-bank/cell-bank-holdings#Humanimmortalmelanocytes). Melanocytes were cultured in RPMI 1640 medium (Lonza) supplemented with 10% fetal bovine serum (FBS), 200 nM TPA (12-O-tetradecanoylphorbol-13-acetate), 10 nM endothelin-1, 200 pM cholera toxin, and 10 ng/mL human stem cell factor. Hermes1 and Hermes3A cells expressing BRAF^V600E^ were cultured without TPA. Melanoma cell lines, including A375 and SKMel28, were purchased from ATCC (Manassas, VA, USA). Additional melanoma cell lines (WM3682, WM164, WM35, WM793, 1205Lu, WM115, WM266.4, and 451Lu) were a gift from Dr. Meenhard Herlyn (Wistar Institute, Philadelphia, PA, USA) from the Wistar Collection of Melanoma Cell Lines. SbCl2 cells were provided by Dr. David Tuveson (Cold Spring Harbor Laboratory, Cold Spring Harbor, NY, USA). Melanocytes were maintained at 37°C in a humidified atmosphere containing 10% CO₂, while melanoma cells were cultured at 37°C with 5% CO₂. HEK293T Lenti-X cells (obtained from Takara) were cultured in DMEM (VWR) containing 10% FBS. All cell lines were routinely tested for mycoplasma contamination using the MycoAlert Mycoplasma Detection Kit (Lonza), and the human melanoma cell lines were STR-authenticated by Moffitt’s Molecular Genomics Core. To inhibit the MAPK pathway, cells were treated with 1 μM Vemurafenib (PLX720, a BRAF inhibitor) or Selumetinib (AZD6244, a MEK1/2 inhibitor) for 48 hours. For transcription and translation inhibition studies, cells were treated with 5 μg/mL actinomycin D or 50 μg/mL cycloheximide, respectively, and harvested at different time points. Golgi inhibitors (Golgi-Stop and Golgi-Plug) were used for 4 hours to inhibit protein transport.

### Plasmids

The plasmids pLenti-CDH3-AS1-Blast and pLenti-CDH3-Hygro were constructed by replacing the GFP coding sequence in pLenti-GFP-Blast and pLenti-GFP-Hygro with CDH3-AS1 (without its intron) or the CDH3 coding sequence (CDS), respectively. CDH3 sequences with or without the 5’UTR were amplified from cDNA. For the generation of the CDH3 construct containing intron 1, the CDH3 intron 1 sequence was amplified from genomic DNA (gDNA) and inserted into the pLenti-CDH3-Hygro plasmid containing the 5’UTR. All cloning steps were performed using the In- Fusion HD Cloning Kit (Takara, Cat. #639650). To add a FLAG tag to the CDH3 sequence, site-directed mutagenesis was performed using the Q5 Site-Directed Mutagenesis Kit (NEB, Ipswich, MA, USA, Cat. #E0554S). The plasmids pLenti-CRISPRa and pLenti-CRISPRv2-puro were obtained from Addgene (Watertown, MA, USA, plasmid #96917 and #52961, respectively). The Cas9 coding sequence was removed from pLenti-CRISPRv2-puro to create a vector exclusively for small guide RNA (sgRNA) expression. Modifications of the sgRNA sequences were performed using the Q5 Site-Directed Mutagenesis Kit according to the manufacturer’s instructions. The psiCHECK2 plasmid was used to construct the double-luciferase reporter plasmids. CDH3 sequences, including the 5’UTR, exon 1 to exon 3, or both, were amplified from gDNA or cDNA and inserted into the double-luciferase vector using the In-Fusion cloning. To generate exon 1 to exon 3 constructs lacking intron 1, site-directed mutagenesis was performed using the Q5 Site-Directed Mutagenesis Kit. All primers used for sequence amplification and site-directed mutagenesis are listed in **Supplementary Table 1**.

### Lentiviral Transduction and Cell Transfection

Lentiviral particles were produced by transfecting HEK293T cells with the lentiviral vector, Δ8.2, and pMD2-VSV-G helper plasmids at a 9:8:1 ratio using JetPRIME transfection reagent (Polyplus, Cat. #101000046). The supernatant from HEK293T cells was collected 48 hours post-transfection, cleared through a 0.45 μm syringe filter, and used to infect target cells in the presence of 8 μg/mL polybrene. Infected cells were selected with 100 µg/mL hygromycin (7 days), 10 µg/mL blasticidin (5 days), or 1 µg/mL puromycin (2 days), depending on the selection marker of the lentiviral construct. Knockdown of *CDH3-AS1* or *CDH3* was achieved using siRNAs. Cells were seeded at 50% confluency and transfected with 25 nM to 50 nM of Dharmacon siRNAs (**see Supplementary Table 1**) using JetPRIME transfection reagent, following the manufacturer’s instructions. JetPRIME was also used for plasmid DNA transfections.

### RNA Sequencing

RNA extracted from Hermes4B cells was quantified with a Qubit Fluorometer (ThermoFisher Scientific, Waltham, MA) and screened for quality on the Agilent TapeStation 4200 (Agilent Technologies, Santa Clara, CA). The samples were then processed for RNA sequencing using the NuGEN Universal RNA-Seq Library Preparation Kit with NuQuant (Tecan Genomics, Redwood City, CA). Briefly, 100 ng of RNA were used to generate cDNA and a strand-specific library following the manufacturer’s protocol. Quality control steps were performed, including TapeStation size assessment and quantification using the Kapa Library Quantification Kit (Roche, Wilmington, MA). The final libraries were normalized, denatured, and sequenced on the Illumina NovaSeq 6000 sequencer with the S1-200 reagent kit to generate approximately 60 million 100- base read pairs per sample (Illumina, Inc., San Diego, CA). Read adapters were detected using BBMerge (v37.02) (19) and subsequently removed with cutadapt (v1.8.1) (20). Processed raw reads were then aligned to human genome GRCh38 using STAR (v2.7.7a) (21). Gene expression was evaluated as read count at gene level with RSEM (v1.3.0) (22) and Gencode gene model v30. Gene express data were then normalized and differential expression between experimental groups was evaluated using DEseq2 (23). Raw counts were used for analyses with BigOmics (https://bigomics.ch/) for the heatmap and GSEA (https://www.gsea-msigdb.org/gsea/index.jsp) analyses for enrichment scores.

### Ribosome profiling (Ribo-seq)

Hermes4B cells were harvested 72 hours after siRNA transfection, and ribosome profiling was performed as described by Cao et al. (24). Briefly, cells grown in 10 cm dishes were washed with 5 mL of ice-cold PBS containing 100 μg/mL cycloheximide (Millipore, Cat. #239764) and immediately harvested by scraping. The cells were collected by centrifugation at 5,000 × g for 5 minutes, and the pellet was resuspended in 400 μL of lysis buffer. The lysate was homogenized by vortexing and incubated on ice for 10 minutes. Clarification was performed by centrifugation at 15,000 × g for 10 minutes at 4°C, and the supernatant was kept on ice. For RNA digestion, 40-60 μg of RNA from the lysate were treated with RNAse I (Lucigen, Cat. #N6901K) for 50 minutes at room temperature while rotating. Following digestion, 5 μL of RNasin Plus RNAse inhibitor (Promega, Cat. #PRN2615) was added to the RNA. Monosome isolation was performed with Microspin S-400 HR columns (Cytiva, Cat. #GE27-5140-01). Columns were washed six times with 500 μL of column wash buffer, followed by centrifugation at 600 × g for 4 minutes.

Subsequently, 100 μL of digested sample was applied to the resin in each column and centrifuged at 600 × g for 2 minutes. Approximately 125 μL of eluate was collected per column, and eluates from the same sample were pooled. To precipitate Ribosome-Protected Fragment (RPF) RNA, an equal volume of acid phenol:chloroform was added to the eluate, vortexed, and incubated at room temperature for 5 minutes. The mixture was centrifuged at 15,000 × g for 10 minutes at 4°C. The aqueous phase was collected, and 1:10 volume of 3 M NaOAc (pH 5.2), 2 μL of glycogen, and 1 volume of ice-cold isopropanol were added. The RPF RNA was precipitated overnight at −80°C. The samples were centrifuged at 15,000 × g for 15 minutes at 4°C, the supernatant was discarded, and the pellet was washed with 80% ethanol. Following another centrifugation at 15,000 × g for 10 minutes at 4°C, the pellet was air-dried and resuspended in 15 μL of nuclease-free water.

To select for the appropriate size RNA, the samples were run on a Novex 12% TBE Urea precast gel (Thermo Fisher Scientific). 5 µL of Gel Loading Buffer (Thermo Fisher, Cat # AM8547) was added to the resuspended RNA and loaded into one or two wells of the gel alongside a 30- nucleotide RNA oligo marker and small RNA ladder (Zymo Cat # R1090). The gel was then washed with TBE, stained with Sybr Gold Nucleic Acid Stain (Thermo Fisher, Cat. #S11494), and imaged. RNA species of 30nt using the marker as reference were excised, crushed, and suspended in 400 μL RNA Binding Buffer (Zymo RNA Clean & Concentrator Kit, Cat. # R1016) and left to diffuse at room temperature overnight. Following overnight diffusion, samples were centrifuged at 10,000 x g for 10 min to pellet the gel and ∼350 μL of RNA binding buffer was removed. The RNA was then isolated using the RNA Clean & Concentrator column (Zymo Research), following the manufacturer’s instructions and eluted in 30 μL of nuclease free water. Following elution, samples were transferred to PCR tubes, mixed with 1 μL (10 units) of T4 Polynucleotide Kinase (NEB, Cat #M0201L), 5 μL T4 Polynucleotide Kinase Reaction Buffer (NEB, Cat #B0201S) and 5 μL 10 mM ATP, and the volume was brought up to 50 μL. Samples were then incubated at 37°C for 1 hour. A modified protocol for the RNA Clean & Concentrator column was then used to isolate RNA. 100 μL adjusted binding buffer (1:1 RNA Binding buffer and ethanol) was mixed with the phosphorylated RNA and added to the column. After centrifugation, 150 μL of ethanol was added to the flow through and this solution was loaded onto a new column and washed according to the manufacturer’s instructions, eluting in 7 μL of nuclease free water. Library preparation was performed using the Takara SMARTer® smRNA-Seq Kit for Illumina® (Takara). Samples received 17 PCR cycles and were sequenced at 50-bp single end reads on a NovaSeq 6000.

### RiboSeq Analysis

Fastq files from RiboSeq libraries were trimmed to remove the first three 5’ nucleotides and all sequences downstream of polyA tracts. Trimmed RPF sequences were then analyzed using RiboToolKit with removal of duplicates and retaining RPFs of 26-34 nt in length. Tri-nucleotide periodicity was analyzed using the RiboWaltz package in R. Differential translation analysis was carried out with DESeq2. For global analysis of translation of antisense containing genes, a curated list of asRNAs was used to categorize transcripts according to the location of the asRNA overlap. RPKM values for transcripts were calculated from the RiboSeq data and analyzed with R. Statistical analysis was performed with Wilcoxon Rank Sum testing.

### Slow Aggregation Assay

96-well plates were pre-filled with 50 μL of FBS-free medium containing 0.1% agarose in each well. After the agarose solidified, 200 μL of cell suspension was added to each well. Images of the resulting spheres were taken at 24 and 48 hours. Cell seeding densities varied depending on the cell line and experimental conditions: SKMel28 cells were plated at 20,000 cells/well, while WM3682, WM793 and Hermes4B cells were plated at 40,000 cells/well. Pictures were taken with EVOS or Incucyte cell imaging system.

### Focus Formation Assay

Cells were seeded in 6-well plates at low density (1,000–2,000 cells/well) and cultured for 14–25 days. Colonies were fixed with methanol and stained for 20 minutes with a 0.1% crystal violet (VWR, Cat# 97061-850) solution prepared in 20% methanol. An equal volume of 10% acetic acid was then added to each well to dissolve the crystal violet. The absorbance of the solution was measured at 600 nm to quantify colony formation.

### RT-PCR and Quantitative RT-PCR

Total RNA was isolated using TRIzol™ Reagent (Invitrogen) following the manufacturer’s protocol. A total of 500 ng RNA per sample was reverse transcribed into cDNA using the PrimeScript RT Master Mix (TakaraBio, Cat. # RR036A). For PCR, cDNA samples were diluted 1:10, and 1 μL of the diluted cDNA was used for amplification. PCR reactions were performed using the Platinum SuperFi II Green PCR Master Mix (Invitrogen) according to the manufacturer’s instructions, with 28 amplification cycles. PCR primers are listed in **Supplementary Table 1**. For quantitative RT-PCR (RT-qPCR), cDNA samples were further diluted at 1:20, and reactions were carried out using the 2X Universal SYBR Green Fast qPCR Mix (ABclonal, Cat # RK21203). Each sample was analyzed in duplicates or triplicates using the StepOne Plus PCR system (Applied Biosystems, Foster City, CA, USA). Relative expression levels were calculated using the comparative threshold cycle method (2^-ΔΔCt), with *GAPDH* serving as the reference transcript for normalization. RT-qPCR primer sequences are listed in **Supplementary Table 1**.

### Cell Fractionation

Cell fractionation was performed using the NE-PER Nuclear and Cytoplasmic Extraction Reagents (Thermo Scientific, #7833) according to the manufacturer’s instructions. Briefly, cell pellets were resuspended in an appropriate volume of ice-cold Cytoplasmic Extraction Reagent I (CER I), vortexed, and incubated on ice for 10 minutes. Cytoplasmic Extraction Reagent II (CER II) was then added, and the mixture was centrifuged at 16,000 × g for 5 minutes at 4°C. The supernatant, corresponding to the cytoplasmic fraction, was collected in fresh tubes. The remaining pellet was resuspended in ice-cold Nuclear Extraction Reagent (NER) and incubated on ice for 40 minutes with intermittent vortexing every 10 minutes. Following centrifugation at 16,000 × g for 10 minutes at 4°C, the supernatant, corresponding to the nuclear fraction, was transferred to a new tube.

RNA from each fraction was extracted using TRIzol™ Reagent (Invitrogen) followed by purification with RNA Clean & Concentrator columns, including DNase treatment. An equal volume of RNA from each fraction was used for reverse transcription. Quantitative PCR (qPCR) data were analyzed using the formula nuclear ratio = 2^-Ct(nuclear)^/2^-Ct(nuclear)^+2^-Ct(cytoplasm)^ and cytoplasmic ratio = 2^-Ct(cytoplasm)^/2^-Ct(nuclear)^+2^-Ct(cytoplasm)^. For western blot analysis, equal volumes of each fraction were mixed with Laemmli buffer (1X final concentration), boiled at 95°C, and subjected to SDS-PAGE.

### RNA Pull-Down with Biotinylated Probes

The RNA pull-down assay was performed following the protocol described by Torres et al. (25). Briefly, cultured cells were washed with PBS and crosslinked using a UV crosslinker set to 150 mJ/cm². Cells were collected and lysed using lysis buffer supplemented with 1% protease inhibitor and RNasin Plus Ribonuclease Inhibitor at 40 U/mL. The lysates were incubated on ice for 15 minutes with vortexing every 5 minutes, followed by sonication for 1 minute. The samples were then centrifuged at 12,000 × g for 5 minutes at 4°C and supernatants were transferred to new tubes. To pre-clear the lysates, streptavidin magnetic beads were added and incubated at room temperature for 30 minutes with rotation. The beads were separated from the lysates, and 20 μL of each lysate was collected as input and stored at −20°C. For hybridization, two volumes of hybridization buffer were added to the lysates, followed by the addition of 100 pmol of biotinylated oligonucleotide probes (see **Supplementary Table 1**). The samples were incubated for 4-6 hours at room temperature with rotation. Pre-washed streptavidin magnetic beads (50 μL) were then added and incubated overnight at 4°C with rotation. After incubation, the beads were separated from the lysates, and the supernatants were discarded. The beads were washed five times with 900 μL of wash buffer. After the final wash, the beads and input samples were resuspended in proteinase K buffer (95 μL for input samples and 75 μL for pull-down samples) and incubated for 45 minutes at 50°C, followed by 10 minutes at 95°C to reverse the crosslinks. The samples were chilled on ice for 3 minutes, and RNA was isolated using RNA Clean & Concentrator columns with DNase treatment, following the manufacturer’s instructions. Reverse transcription was performed using an equal volume of RNA for each sample condition. qPCR data were analyzed and normalized to the input samples.

### Immunoblotting

Protein lysates were prepared from cell pellets or mouse tumor tissues. Samples were homogenized in RIPA buffer (50 mM Tris-HCl, pH 8.0, 1 mM EDTA, 0.5 mM EGTA, 1% Triton X- 100, 0.5% sodium deoxycholate, 0.1% SDS, 150 mM NaCl) supplemented with protease and phosphatase inhibitors. Total protein (20 μg per sample) was mixed with Laemmli buffer, boiled, and separated on NuPAGE 4–12% precast gels (Thermo Fisher Scientific). Proteins were transferred to nitrocellulose membranes using standard wet-transfer protocols. Membranes were blocked in 5% non-fat dry milk prepared in TBST (20 mM Tris, 150 mM NaCl, 0.1% Tween-20) for 1 hour at room temperature and incubated overnight at 4°C with primary antibodies diluted in 5% milk or BSA. The following primary antibodies were used: P-cadherin (BD Bioscience, #610227; 1:1,000), ERK1/2 (CST, #4695; 1:5,000), phospho-ERK1/2 (Thr202/Tyr204, CST, #9101; 1:2,000), Laminin A/C (CST, #2032S; 1:1,000), α-Tubulin (DM1A, CST, #3873S; 1:2,000), MCL1 (Santa Cruz Biotechnology, #sc-12756; 1:5,000), FLAG (M2, Millipore, #F1804; 1:1,000), GFP (CST, #2956S; 1:2,000), and β-Actin (Invitrogen, AM4302; 1:10,000), which served as the loading control. Membranes were washed four times for 10 minutes each in TBST, followed by incubation with horseradish peroxidase-conjugated secondary antibodies (1:10,000) for 1 hour at room temperature. After four additional washes in TBST, the blots were developed using chemiluminescent substrate (1:1 dilution) for 3 minutes. Signals were visualized using a LI-COR imaging system or ProSignal Blotting Film (Prometheus).

### Animal Experiments

Animal experiments were conducted in accordance with an Institutional Animal Care and Use Committee protocol approved by the University of South Florida (Tampa, FL). NSG mice (stock no. 005557) were obtained from The Jackson Laboratory (JAX) and bred in-house under pathogen-free conditions. For xenograft experiments, mice were anesthetized with isoflurane, and the dorsal area was shaved using clippers. A total of 5 × 10^5^ cells in a single-cell suspension were injected subcutaneously into the flanks of female NSG mice. Tumor growth was monitored using digital calipers, and tumor volume was calculated using the formula: (width^2^ x length)/2.

### Quantification and statistical analysis

Data are presented as the mean ± standard error of the mean (SEM). All experiments were performed at least three times with two to five technical replicates and one representative experiment is shown. Statistical comparisons between two groups were performed using an unpaired two-tailed t-test. A p-value of < 0.05 was considered statistically significant.

## RESULTS

### The lncRNA expression landscape in melanoma

To comprehensively analyze lncRNA expression changes during melanocyte transformation and melanoma formation, we analyzed RNA sequencing data from four human melanocyte cell lines (Hermes1, Hermes2, Hermes3A, Hermes4B) and five human melanoma cell lines (WM35, WM793, WM164, SKMel28, 1205Lu) (**Figure 1A, GSE148552**). This analysis revealed 8,984 differentially expressed genes (adj p-value < 0.05; −1 > Log2(FC) > 1), with a significant portion being ncRNAs (43.6%). Among these, three principal groups represent the majority of the non-coding transcriptome: pseudogenes, lincRNAs, and asRNAs (**Figure 1A, Supplementary Table 2)**. asRNAs were the third most deregulated group of ncRNAs in melanoma, and due to their overlap with sense genes, they have significant regulatory potential. Differentially expressed asRNA genes predominantly overlap with a single sense gene (**Figure 1B**). However, in some cases, asRNA genes overlap with multiple sense genes, with one antisense transcript overlapping five sense genes, while other genes classified as asRNA genes do not overlap with known protein-coding genes (**Figure 1B**). We found that 73.4% of deregulated asRNA genes overlap with a single sense gene, while those overlapping with two or more sense genes represent 13.6% of the deregulated asRNAs (**Figure 1B**).

**Figure 1:**
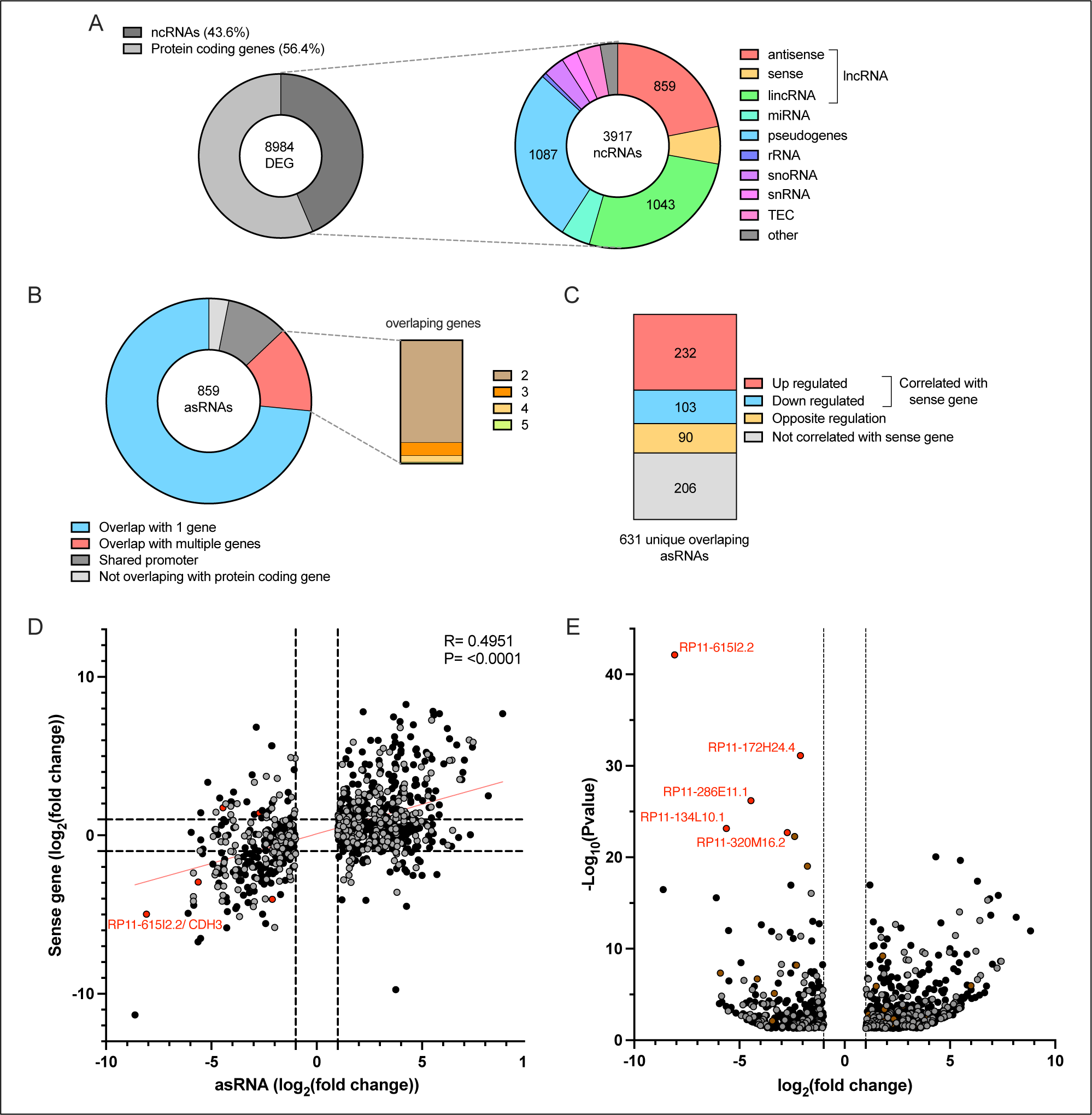
The asRNA landscape of melanoma. (A) Biotypes of differentially expressed genes in melanoma cell lines. (B) Genomic position of asRNA genes relative to their corresponding sense protein coding genes and number of overlapping protein coding sense genes. (C) Distribution of sense/antisense gene pairs with concordant or discordant expression patterns. (D) Fold-change expression of deregulated asRNAs and their corresponding sense protein coding genes in melanoma. Black dots represent asRNAs overlapping a single sense protein-coding gene, red dots represent the most downregulated asRNA, gray dots represent asRNAs overlapping multiple genes. (E) Volcano plot showing differentially expressed asRNAs in melanoma compared to melanocytes. Brown dots represent asRNAs overlapping a single sense non-coding RNA gene not listed as protein coding gene.

While some asRNA genes were differentially expressed in the absence of expression changes of the corresponding sense gene, more than half of the asRNA genes overlapping with a single sense gene exhibited expression concordance (**Figure 1C,D**). This suggests either transcriptional co-regulation of antisense/sense gene pairs or an impact of asRNAs on their cognate sense mRNAs. We next aimed to select asRNAs that are strongly deregulated in melanoma for functional studies. Differential gene expression analysis of annotated asRNAs revealed that the top five deregulated asRNAs are significantly downregulated in melanoma (**Figure 1E)**. Among these, we focused on *CDH3-AS1* (annotated as RP11-615I2.2) because it was the most downregulated asRNA in melanoma compared to melanocytes and the expression of its cognate sense protein-coding gene, *CDH3*, was also significantly reduced (**Figure 1D**). These observations suggested the involvement of asRNAs in melanoma formation and prompted further investigation into the role of *CDH3-AS1* regulation in melanoma.

### MAPK pathway activation reduces *CDH3-AS1* and *CDH3* expression in melanoma

*CDH3-AS1* is a 1,682 nucleotide asRNA composed of two exons and is classified as a head-to-head asRNA (12, 26). It overlaps with exons 1 and 2 of the *CDH3* protein-coding gene (**Figure 2A, Supplementary Figure 1A**). *CDH3* encodes P-cadherin, a transmembrane protein belonging to the cadherin superfamily, which plays a critical role in calcium-dependent cell-cell adhesion (27, 28). To validate the downregulation of the *CDH3-AS1*/*CDH3* pair in melanoma, we performed RT-qPCR on a panel of human melanocyte and melanoma cell lines. This analysis confirmed our RNA sequencing results, showing that both *CDH3-AS1* and *CDH3* were expressed in melanocytes but were undetectable in all but two of the melanoma cell lines analyzed (**Figure 2B**). Consistently, *CDH3*-encoded P-cadherin protein levels mirrored the mRNA downregulation (**Figure 2C**). We also assessed the presence of the 50 kDa isoform of P-cadherin (29) but did not detect its expression in either melanocytes or melanoma cells (**Supplementary Figure 1B**). We observed that compared to the BRAF wildtype Hermes1 and Hermes3A melanocytes, the BRAF^V600E^-mutant derivative cell lines H1B and H3B (30) exhibited a marked decrease in *CDH3-AS1* and *CDH3* expression (**Figure 2C**). Given the association of the BRAF^V600E^ mutation with reduced expression of *CDH3-AS1* and *CDH3*, we investigated whether the MAPK pathway mediates the downregulation of the *CDH3-AS1*/*CDH3* pair. Hermes melanocytes are cultured in the presence of growth factors, including the phorbol ester TPA that stimulates the MAPK pathway. We therefore tested whether TPA impacted *CDH3-AS1*/*CDH3* expression in Hermes melanocytes. TPA withdrawal from the media of Hermes1 and Hermes4B melanocytes led to a reduction in pERK levels, indicating decreased MAPK pathway activity, and resulted in a significant increase in *CDH3-AS1* and *CDH3* mRNA and P-cadherin protein levels (**Figure 2D,E**).

**Figure 2:**
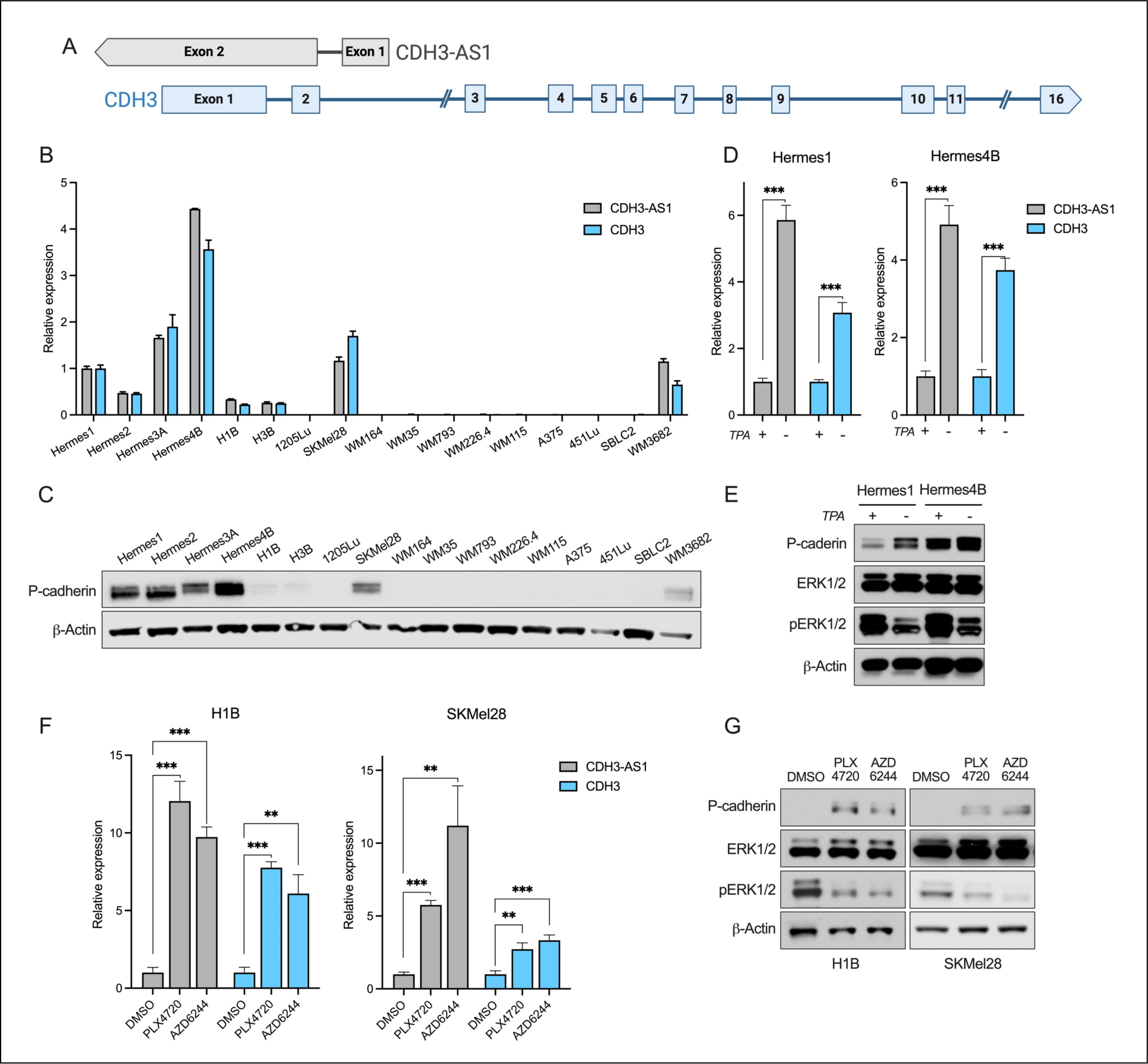
The MAPK pathway downregulates CDH3-AS1 and CDH3 in melanoma. (A) Schematic representation of the overlapping CDH3-AS1 and CDH3 genes in a head-to-head orientation. (B-C) Relative expression levels of CDH3-AS1 and CDH3 measured by RT-qPCR (B) and P-cadherin protein levels analyzed by Western blot (C) in human melanocytes and melanoma cell lines. (D-E) Relative expression levels of CDH3-AS1 and CDH3 (D) and Western blot analysis of P-cadherin protein levels (E) in Hermes1 and Hermes4B melanocytes cultured with or without TPA for 48 hours. (F-G) RT-qPCR analysis of CDH3-AS1 and CDH3 RNA levels (F) and Western blot analysis of P-cadherin protein levels (G) in H1B and SKMel28 cells treated with PLX4720 (BRAF inhibitor) or AZD6244 (MEK1/2 inhibitor) for 48 hours. * = p < 0.05; ** = p < 0.01; *** = p < 0.001.

To further validate these findings, we inhibited the MAPK pathway using the BRAF inhibitor Vemurafenib (PLX4720) or the MEK inhibitor Selumetinib (AZD6244) in BRAF^V600E^-mutant melanocytes and melanoma cells expressing low baseline levels of *CDH3-AS1* and *CDH3*. Both H1B and SKMel28 cells showed a significant increase in *CDH3-AS1* and *CDH3* mRNA levels and P-cadherin protein levels when the MAPK pathway was inhibited (**Figure 2F,G**). These results indicate that *CDH3-AS1* and *CDH3* are strongly downregulated in melanocytes and melanoma cells upon activation of the MAPK pathway.

#### *CDH3-AS1* regulates P-cadherin translation preferentially *in cis*

We investigated whether *CDH3-AS1* and *CDH3* are independently downregulated in melanoma cells or if their expression is interdependent. Given that *CDH3-AS1* is antisense to *CDH3* and the two RNAs harbor complementarity, we hypothesized that *CDH3-AS1* regulates *CDH3* expression. To test this, we increased *CDH3-AS1* expression using two strategies: CRISPR activation (CRISPRa) to examine cis-regulation and lentiviral overexpression to study trans-regulation. Both methods successfully upregulated *CDH3-AS1* but had minimal impact on *CDH3* mRNA levels (**Figure 3A**). Interestingly, CRISPRa led to a 6-fold increase in *CDH3-AS1* and an 18-fold increase in P-cadherin levels (**Figure 3A,B**). In contrast, lentiviral overexpression of *CDH3-AS1*, resulting in a nearly 400-fold increase in asRNA levels, caused only a modest rise in P-cadherin expression (**Figure 3A,B**). To further explore this regulation, we co-transfected HEK293T cells with equal amounts of *CDH3* (full exonic sequence) and increasing amounts of *CDH3-AS1*. While *CDH3* mRNA levels remained unaffected, P-cadherin levels increased proportionally with higher *CDH3- AS1* expression (**Figure 3C,D**). These findings suggest that *CDH3-AS1* enhances P-cadherin levels without directly affecting *CDH3* mRNA abundance and that this regulation is more efficient in cis, when *CDH3-AS1* and *CDH3* are expressed from a common locus.

**Figure 3:**
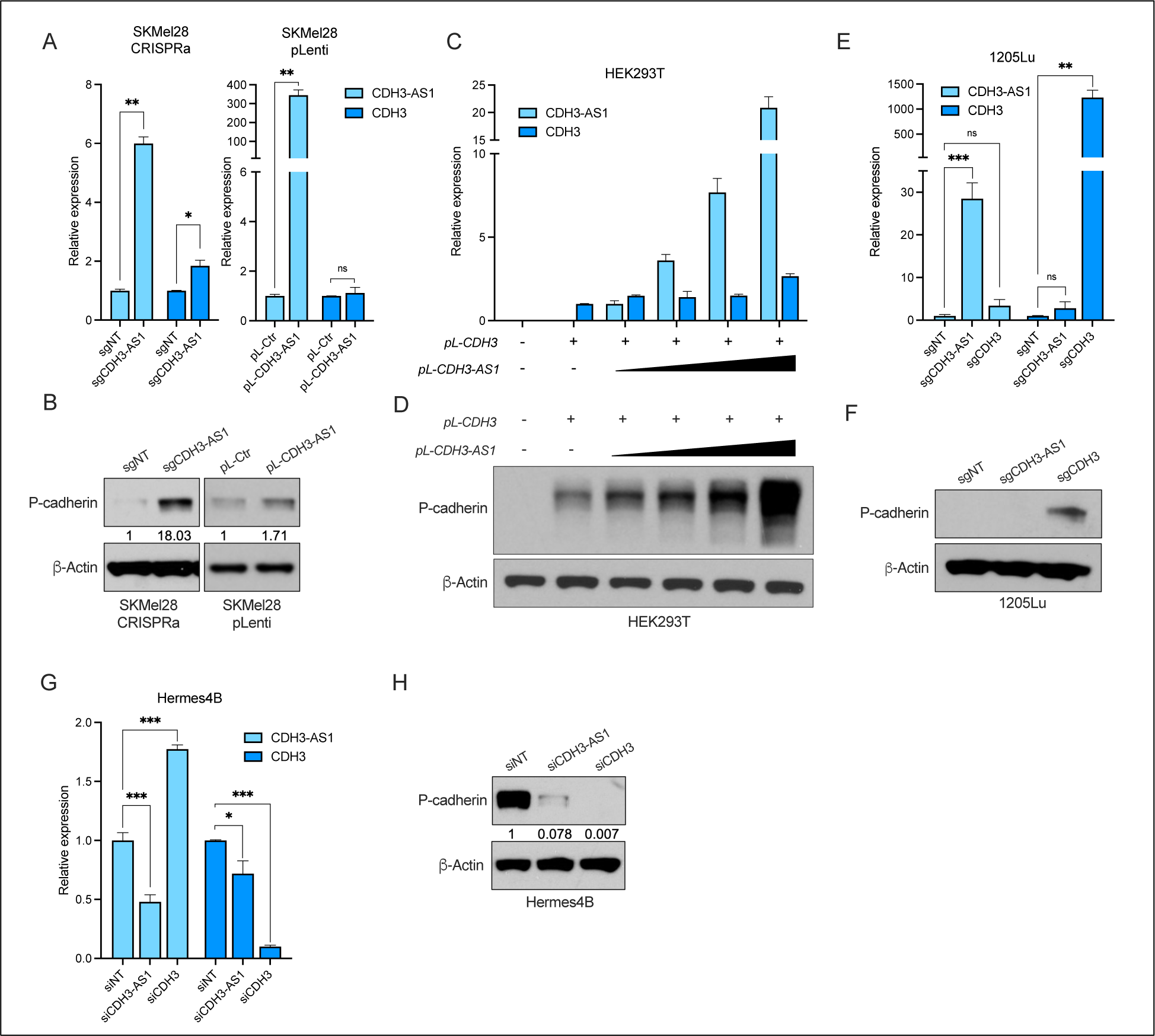
CDH3-AS1 increases P-cadherin protein levels. (A-B) RT-qPCR analysis of CDH3-AS1 and CDH3 expression (A) and Western blot analysis of P- cadherin protein levels (B) in SKMel28 cells following CRISPRa- or lentiviral-mediated CDH3- AS1 overexpression. (C-D) RT-qPCR analysis of CDH3-AS1 and CDH3 expression (C) and Western blot showing P-cadherin protein levels (D) in HEK293T cells co-transfected with full- length pL-CDH3 plasmid and increasing amounts of pL-CDH3-AS1 plasmid. (E-F) RT-qPCR analysis of CDH3-AS1 or CDH3 expression (E) and Western blot analysis of P-cadherin levels (F) in 1205Lu cells. (G-H) RT-qPCR analysis of CDH3-AS1 and CDH3 expression (G) and Western blot analysis of P-Cadherin protein levels (H) in Hermes4B melanocytes following knockdown of CDH3-AS1 or CDH3 using siRNAs. ns = not significant; * = p < 0.05; ** = p < 0.01; *** = p < 0.001.

SKMel28 cells exhibit low baseline expression of both *CDH3-AS1* and *CDH3*, whereas other melanoma cell lines, such as 1205Lu, show no baseline expression (**Figure 2B**; **Figure 3E,F**). We used a CRISPRa strategy to determine if overexpression of *CDH3-AS1* could elevate P- cadherin protein levels in 1205Lu cells. While increasing *CDH3* expression led to increased P- cadherin levels, P-cadherin remained undetectable upon *CDH3-AS1* overexpression despite a 30-fold increase (**Figure 3E,F**). These findings indicate that baseline *CDH3* expression is required for *CDH3-AS1* to modulate P-cadherin levels.

To further validate the regulation of P-cadherin by *CDH3-AS1*, we silenced *CDH3-AS1* using small interfering RNA (siRNA) in Hermes4B melanocytes and assessed P-cadherin levels. siCDH3 reduced *CDH3* mRNA levels by 90%, leading to a complete loss of P-cadherin protein levels (**Figure 3G,H**). The siRNA against *CDH3-AS1* reduced the asRNA expression levels by 50%, with a concomitant modest decrease (∼20%) in *CDH3* mRNA levels. Interestingly, *CDH3-AS1* silencing resulted in a ∼10-fold decrease in P-cadherin levels (**Figure 3G,H**), which is unlikely to be accounted for by the modest reduction in *CDH3* mRNA levels. Additionally, we observed a more than 1.5-fold increase in *CDH3-AS1* expression upon *CDH3* silencing, suggesting a potential feedback loop between *CDH3-AS1* and P-cadherin. Collectively, these results demonstrate that *CDH3-AS1* regulates P-cadherin levels with only minimal effects on *CDH3* mRNA levels.

### *CDH3-AS1* modulation phenocopies the tumor suppressive effects of P-cadherin

Because *CDH3-AS1* influences P-cadherin protein levels, we hypothesized that overexpression of *CDH3-AS1* mimics the phenotype observed upon increased P-cadherin expression. While the role of P-cadherin has been studied (28), its function in melanoma remains controversial, with reports suggesting both tumor suppressive (31–33) and oncogenic roles (34, 35). For this phenotypic study, we utilized CRISPRa to overexpress *CDH3-AS1* or *CDH3* in SKMel28 and WM3682 melanoma cell lines. This approach successfully increased RNA expression of *CDH3- AS1* or *CDH3*, leading to elevated P-cadherin levels under both conditions in both cell lines (**Supplementary Figure 2A,B**). To assess the functional impact of these changes, we conducted cell biology assays. Interestingly, no significant differences in cell proliferation were observed compared to control cells when either *CDH3* or *CDH3-AS1* was overexpressed (**Supplementary Figure 2C**). In contrast, overexpression of *CDH3* led to a reduction in colony formation in WM3682 cells and *CDH3-AS1* overexpression phenocopied these effects (**Figure 4A**). In SKMel28 cells, *CDH3* overexpression only modestly reduced colony formation and *CDH3-AS1* had no effects (**Supplementary Figure 2D**), suggesting that P-cadherin may not play a consistent and prominent role in cell growth across melanoma cell lines. Given that P-cadherin is a cell adhesion protein that promotes cell aggregation (31), we performed aggregation assays. This revealed that cells overexpressing *CDH3-AS1* formed more compact spheres resulting in smaller sphere sizes, similar to the effects observed with *CDH3* overexpression (**Figure 4B, Supplementary Figure 2E**). Conversely, knockdown of *CDH3-AS1* or *CDH3* increased sphere sizes, indicating reduced cell aggregation (**Figure 4C**). We also conducted cell aggregation assays using WM793 melanoma cells which lack baseline expression of *CDH3* and *CDH3-AS1*. Lentiviral *CDH3* expression resulted in increased P-cadherin levels (**Supplementary Figure 2F**) and reduced sphere sizes (**Figure 4D**). In contrast, *CDH3-AS1* overexpression did not elevate P-cadherin levels (**Supplementary Figure 2F**) and accordingly did not lead to a decrease in sphere size (**Figure 4D**). These findings suggest that *CDH3-AS1* promotes cell aggregation through its regulation of P-cadherin.

**Figure 4:**
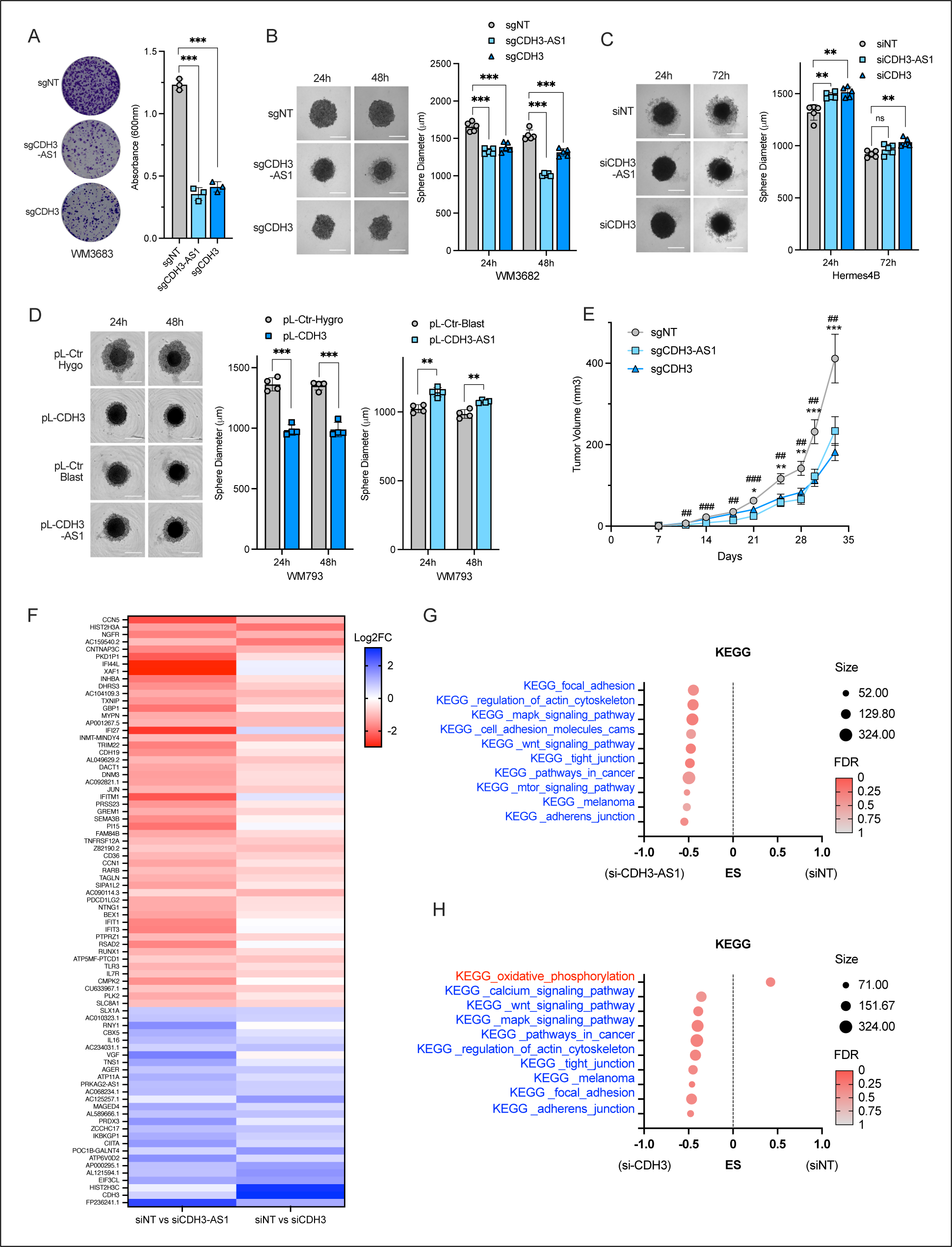
CDH3-AS1 and P-cadherin are tumor suppressors in melanoma. (A) Low density colony formation assay and absorbance measurements in WM3682 cells overexpressing CDH3-AS1 or CDH3. (B-D) Slow aggregation assay showing sphere diameter measurements in WM3682 cells with CRISPRa stable overexpression (B), in Hermes4B cells with transient downregulation (C) or in WM793 cells with lentiviral stable overexpression of CDH3-AS1 or CDH3. (E) Tumor growth curves of WM3682 cells overexpressing CDH3-AS1 or CDH3 transplanted into NSG mice. Two tumors were inoculated per mouse and six mice per group were analyzed. # depicts statistical significance between sgNT and sgCDH3-AS1; * depicts statistical significance between sgNT and sgCDH3. (F) Heatmap showing overlapping expression profiles of deregulated genes in Hermes4 melanocytes comparing siNT vs siCDH3-AS1 with siNT vs siCDH3. (G-H) Gene Set Enrichment Analysis (GSEA) of differentially expressed genes in Hermes4B cells upon silencing of CDH3-AS1 (G) or CDH3 (H), based on KEGG pathways. * or # = p < 0.05; ** or ## = p < 0.01; *** or ### = p < 0.001.

To examine the tumor suppressor function of *CDH3-AS1* in vivo, we analyzed tumor growth of WM3682 CRISPRa cells subcutaneously xenografted in immunocompromised NSG mice. Overexpression of either *CDH3-AS1* or *CDH3* significantly reduced tumor volumes compared to the non-targeting sgRNA control (**Figure 4E**). Protein analysis of the tumors confirmed that *CDH3- AS1* overexpression increased P-cadherin levels (**Supplementary Figure 2G**). These in vitro and in vivo results validate the tumor suppressor role for P-cadherin, demonstrate a tumor suppressor role for *CDH3-AS1*, and suggest that *CDH3-AS1* exerts its tumor suppressive effects through the regulation of P-cadherin levels.

To further assess whether *CDH3-AS1* exerts its tumor suppressor function through the regulation of P-cadherin, we silenced *CDH3-AS1* or *CDH3* in Hermes4B cells and performed RNA sequencing. The silencing of *CDH3-AS1* induced transcriptional changes similar to the silencing of *CDH3* (**Figure 4F**). Moreover, GSEA analyses of differentially expressed genes revealed that silencing *CDH3-AS1* or *CDH3* affects similar KEGG pathways and biological processes, including the MAPK signaling pathway, tight junctions, melanoma, and adherens junctions (**Figure 4G,H**). Additionally, we identified a shared association with the epithelial-to-mesenchymal transition hallmark in the *CDH3-AS1* and *CDH3* knockdown samples (**Supplementary Figure 2H,I**). The similarities in gene expression changes and enriched pathways suggest that *CDH3-AS1* functions through the regulation of P-cadherin.

### *CDH3-AS1* does not influence the splicing, localization, or stability of *CDH3*

To better understand the mechanism whereby *CDH3-AS1* regulates P-cadherin levels, we first determined its cellular localization. *CDH3-AS1* is primarily localized in the cytoplasm of both melanocytes and melanoma cell lines, similar to *CDH3*, which is consistent with the fact that *CDH3* is a protein-coding mRNA (**Figure 5A, Supplementary Figure 3A**). Additionally, we confirmed that P-cadherin is also localized in the cytoplasm fraction containing the plasma membrane, indicating that *CDH3-AS1* does not affect P-cadherin localization (**Supplementary Figure 3B**). Given that both RNAs are localized within the same cellular compartment and that *CDH3-AS1* elicits a stronger effect on P-cadherin levels when transcribed in close proximity to the *CDH3* mRNA, we investigated whether these RNAs interact directly. To test this, we performed RNA pull-down assays using biotinylated probes targeting *CDH3-AS1* exon 2 in Hermes4B cells. This approach readily isolated *CDH3-AS1* (**Supplementary Figure 3C**) and, importantly, resulted in the enrichment of *CDH3* mRNA (**Figure 5B**). *GAPDH* mRNA served as negative control and showed negligible enrichment (**Figure 5B**). To further confirm the interaction between *CDH3-AS1* and *CDH3*, we conducted co-transfection experiments in HEK293 cells using a constant amount of pLenti-CDH3 plasmid and increasing amounts of pLenti-CDH3-AS1 plasmid. Subsequent RNA pull-down assays demonstrated that higher *CDH3-AS1* expression resulted in greater enrichment of *CDH3-AS1* (**Supplementary Figure 3D**), leading to a dose-dependent increase of co-purified *CDH3* (**Figure 5C**). The above pull-down assays used sonication to fragment the RNA and primers in exons 12/13 to detect co-purified *CDH3*. If *CDH3-AS1* interacts with exons 1/2 of *CDH3*, then this protocol may result in inefficient co-purification of *CDH3*. We therefore performed pull-down assays using additional primers in exons 2/3 near the presumed interaction site in *CDH3* (**Figure 5D, Supplementary Figure 3E**). Interestingly, the *CDH3* enrichment with primers in exons 2/3 was 5-fold greater than with primers in exons 12/13, suggesting that *CDH3-AS1* interacts with the 5’ region of *CDH3* (**Figure 5D**).

**Figure 5:**
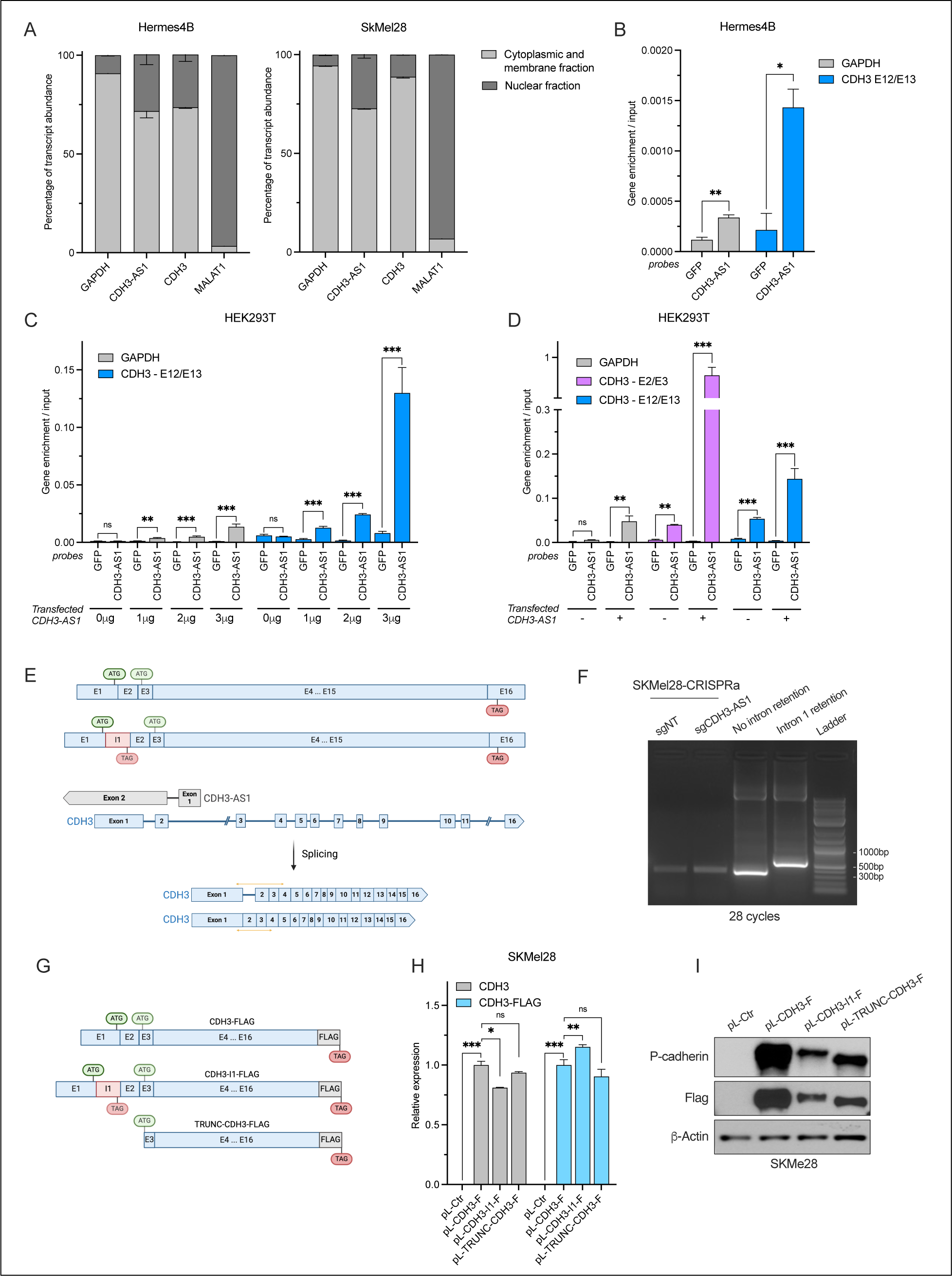
CDH3-AS1 interacts with CDH3 but does not affect its splicing. (A) qPCR showing GAPDH (cytoplasmic control), CDH3-AS1, CDH3, and NEAT1 (nuclear control) transcripts in Hermes4B and SKMel28 cells after cell fractionation to separate nuclear and cytoplasmic/plasma membrane fractions. (B-D) Enrichment of GAPDH or CDH3 mRNA in RNA pull-down assays using biotinylated probes against CDH3-AS1 in Hermes4B cells (B) or in HEK293T cells co-transfected with equal amounts of pL-CDH3 and increasing amounts of pL- CDH3-AS1 plasmids (C), or with primer pairs targeting CDH3 exon2/3 (E2/E3) or exon12/13 (E12/E13) (D). (E) Schematic representation of the CDH3 gene indicating the start codon (ATG) and stop codon (TAG), with a diagram showing potential intron retention. PCR amplicons are indicated by yellow arrows. (F) Agarose gel electrophoresis of PCR-amplified CDH3 cDNA showing intron 1 exclusion (362 bp band) and intron 1 inclusion (573 bp band). (G) Diagram of three CDH3-FLAG (CDH3-F) constructs used for transient transfection experiments. (H, I) qPCR (H) and Western blot (I) showing CDH3 mRNA expression and P-cadherin protein levels, respectively, in SKMel28 cells 24 hours after transient transfection with equal amounts of plasmid DNA. * = p<0.05; ** = p<0.01; *** = p<0.001. (E) and (G) were generated with BioRender. CDH3 E2-E3 or E12-13: primer pair targeting exon2-exon3 or exon12-exon13 of CDH3, respectively.

Having demonstrated the interaction between *CDH3-AS1* and *CDH3*, we next sought to investigate the mechanism by which *CDH3-AS1* influences P-cadherin levels. First, we examined whether *CDH3-AS1* stabilizes the *CDH3* mRNA. To test this, we treated cells with actinomycin D to inhibit transcription and assessed *CDH3* mRNA turnover. Overexpression of *CDH3-AS1* in SKMel28 melanoma cells using pLenti-CDH3-AS1 did not increase the stability of *CDH3* (**Supplementary Figure 3F**) nor did this expression approach affect the stability of *CDH3-AS1* (**Supplementary Figure 3G**). Using cycloheximide, we also demonstrated no changes in P-cadherin stability in response to *CDH3-AS1* overexpression (**Supplementary Figure 3H**). These results suggest that *CDH3-AS1* does not affect the stability of either *CDH3* mRNA or P-cadherin protein.

Previous studies have shown that asRNAs can influence translation through various mechanisms, such as altering RNA splicing of the sense RNA to promote alternative protein isoforms (36), or preserving an IRES within the sense RNA to enhance protein translation (37). Because *CDH3- AS1* overlaps with exons 1 and 2 as well as intron 1 of *CDH3* (**Figure 2A**), we investigated whether *CDH3-AS1* impacts splicing of *CDH3* intron 1. The canonical ATG start codon of *CDH3* is in exon 1 and intron 1 contains a stop codon in-frame with the canonical start codon. Thus, if intron 1 were retained, *CDH3* translation would terminate prematurely unless an alternative ATG start codon downstream of intron 1 and in-frame with the ORF is used to initiate translation. Such an ATG start codon exists at the beginning of exon 3 (**Figure 5E**). We examined whether *CDH3- AS1* promotes *CDH3* intron 1 retention, leading to P-cadherin translation from the downstream ATG to produce a shorter isoform. Using RT-PCR analysis, we observed no retention of intron 1 at baseline or when *CDH3-AS1* was overexpressed in SKMel28 cells (**Figure 5F**). To further confirm these results and determine which *CDH3* start codon is primarily used, we generated three flag-tagged *CDH3* cDNA constructs: one containing the full-length *CDH3* coding sequence, one with intron 1 retained, and a truncated isoform starting from the downstream ATG (**Figure 5G**). Transient transfection of these constructs into SKMel28 cells revealed that all three constructs were expressed at similar mRNA levels (**Figure 5H**), but P-cadherin protein levels varied significantly (**Figure 5I**). The construct containing the full-length *CDH3* coding sequence showed the highest translation efficiency, while the intron 1-containing and truncated constructs exhibited reduced translation efficiency and lower P-cadherin protein levels (**Figure 5I**). Furthermore, the molecular weight of the P-cadherin protein produced from the intron 1-containing *CDH3* construct suggested that the second ATG is not used in the presence of intron 1. Rather, the canonical ATG is used and intron 1 is spliced out, resulting in the translation of full-length P-cadherin. We conclude that *CDH3-AS1* does not promote intron 1 retention to promote the translation of a shorter P-cadherin isoform from an alternative downstream start codon.

It was recently demonstrated that asRNAs can accelerate mRNA nuclear export by annealing with their sense counterparts (38). We therefore investigated whether *CDH3-AS1* influences *CDH3* nuclear export. To this end, we performed cell fractionation on Hermes4B cells treated with siRNA targeting *CDH3-AS1* and assessed the localization of *CDH3* mRNA. We observed no significant changes in *CDH3* mRNA localization upon *CDH3-AS1* silencing (**Supplementary Figure 3I**). However, P-cadherin protein levels were markedly reduced under the same conditions (**Supplementary Figure 3J**). These results indicate that *CDH3-AS1* does not regulate P-cadherin protein levels by promoting the export of *CDH3* mRNA to the cytoplasm.

### *CDH3-AS1* facilitates the translation of P-cadherin

We next investigated whether *CDH3-AS1* influences the translation efficiency of *CDH3* mRNA. We hypothesized that secondary structures in the 5’UTR region of *CDH3* could impair translation and that *CDH3-AS1* facilitates the resolution of such structures. To assess the presence of secondary structures in *CDH3*, we treated Hermes4B and SKMel28 cells with Silvestrol, an inhibitor of the RNA helicase eIF4A that unwinds RNA secondary structures to facilitate pre- initiation complex recruitment and translation (39, 40). Silvestrol treatment resulted in a reduction of P-cadherin protein levels without a corresponding change in *CDH3* mRNA levels (**Figure 6A, Supplementary Figure 4A**). These findings suggest that *CDH3* mRNA harbors a secondary structure that impacts its translation. Interestingly, Silvestrol treatment also increased *CDH3-AS1* RNA levels, suggesting a possible MAPK-independent feedback loop in which *CDH3-AS1* is upregulated when P-cadherin levels decrease (**Supplementary Figure 4A**).

**Figure 6:**
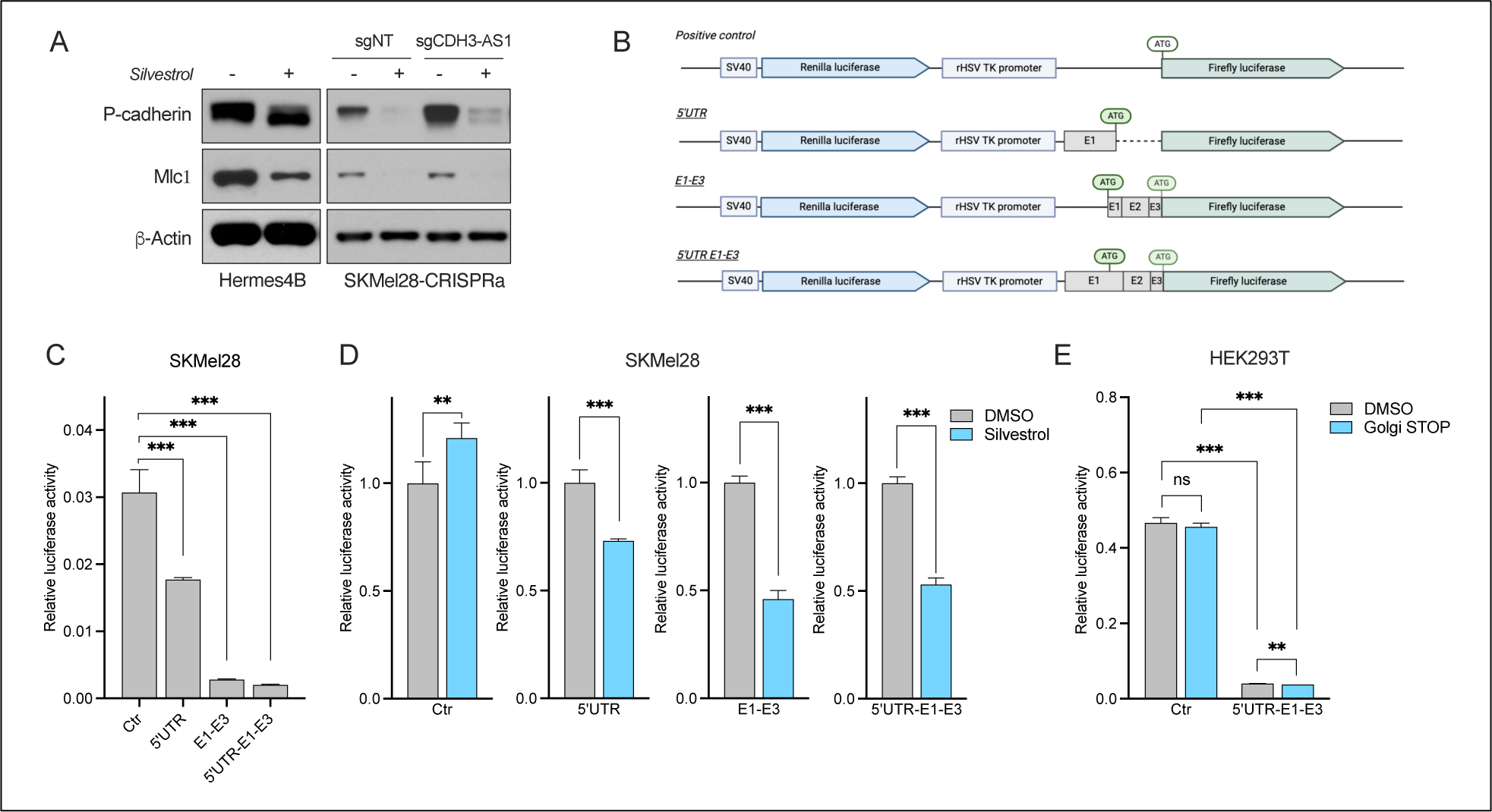
A secondary structure in the 5’end of CDH3 impairs P-cadherin translation. (A) P-cadherin protein levels in Hermes4B and SKMel28 cells following treatment with Silvestrol for 24 hours. (B) Schematic representation of four p-Lenti dual luciferase constructs used for transfection and luciferase activity assays. Made with BioRender. (C) Relative luciferase activity of the four constructs shown in (B) in SKMel28 cells. (D) Relative luciferase activity of the four constructs following 24 hours of Silvestrol treatment in SKMel28. (E) Relative luciferase activity in HEK293T cells co-transfected with equal amounts of dual luciferase constructs and treated with DMSO or Golgi STOP mix. * = p<0.05; ** = p<0.01; *** = p<0.001; ns = not significant. Given that both the CDH3 5’UTR and early CDS impair P-cadherin translation, we tested the effect of CDH3-AS1 on the 5’UTR of CDH3. To this end, we transfected SKMel28 cells with pLenti- CDH3 constructs containing either the 5’UTR region or only the CDS. While CDH3 mRNA levels remained unchanged (**Supplementary Figure 4B**), P-cadherin protein levels were reduced when CDH3 mRNA included the 5’UTR (**Supplementary Figure 4C**). Notably, CDH3-AS1 overexpression increased P-cadherin protein levels also when the 5’UTR was present (**Supplementary Figure 4C**). Overall, these findings suggest that CDH3-AS1 influences secondary structure elements in the 5’UTR and early CDS of CDH3 to facilitate P-cadherin translation.

Sensitivity to eIF4A inhibition has been shown to correlate with the complexity of secondary structures in the 5’UTR (39). Since *CDH3-AS1* overlaps *CDH3* from the 5’ UTR to exon 2 (**Figure 2A**), we examined whether this region contains secondary structures impacting translation. We created several double-luciferase reporter constructs containing either the *CDH3* 5’UTR alone, the coding sequence (CDS) from the canonical ATG in exon 1 to the second ATG in exon 3 (E1- E3), or the complete 5’UTR to exon 3 region (5’E1-E3) (**Figure 6B**). The luciferase activity decreased by approximately 50% in the 5’UTR construct and by around 90% in both the E1-E3 and 5’E1-E3 constructs (**Figure 6C**). Treatment with Silvestrol further reduced the luciferase activity of the 5’UTR, E1-E3 and 5’E1-E3 constructs while the control luciferase activity was resistant to Silvestrol, suggesting the presence of secondary structures in the 5’UTR and early CDS regions of *CDH3* mRNA that impair translation (**Figure 6D**). To validate that the decrease in luciferase activity is due to secondary structures rather than P-cadherin signal peptide-mediated transport of the luciferase protein to the membrane for secretion, we treated HEK293T cells with Golgi Stop and Golgi-Plug for 4 hours. The treatment did not affect luciferase activity in the control and did not rescue the reduction of luciferase activity of the 5’E1-E3 construct (**Figure 6E**). This indicates that the reduced luciferase activity is due to secondary structures in this region.

### *CDH3-AS1* influences ribosome binding to *CDH3*

To gain deeper insight into how *CDH3-AS1* influences *CDH3* translation, we performed ribosome profiling on Hermes4B cells to map ribosome distribution across *CDH3* mRNA at baseline and in response to *CDH3-AS1* silencing. After size selection of ribosome-protected RNA fragments (RPFs, **Supplementary Figure 5A**), we verified that the length of sequenced RPFs was consistent across samples (**Supplementary Figure 5B**). Global translational activity remained unperturbed upon *CDH3-AS1* knockdown, as evidenced by robust capture of translating ribosomes in both siNT and siCDH3-AS1 conditions (41) (**Figure 7A, Supplementary Figure 5C**). Integrated analysis of differential mRNA abundance and ribosome occupancy identified *CDH3* as one of the most differentially translated genes upon *CDH3-AS1* downregulation (**Figure 7B**). High-resolution ribosome profiling further revealed depleted ribosome occupancy in both the 5’UTR and CDS of *CDH3* in the absence of *CDH3-AS1* (**Figure 7C**), while mapped P-sites were consistent and aligned in the same reading frame between conditions (**Supplementary Figure 5D,E**). Collectively, these results suggest that *CDH3-AS1* modulates *CDH3* mRNA translation. We then investigated whether ribosome occupancy of sense protein-coding transcripts is influenced by the presence of asRNAs. Using the ribosome profiling dataset, we identified transcripts with asRNA and determined the regions of overlap for each transcript (**Supplementary Table 3**). Interestingly, transcripts having asRNA whose complementarity covers the 5’ region, extending from the 5’UTR to the CDS similar to the structure of *CDH3-AS1*, exhibited a significant increase in translation (**Figure 7D**). We then considered the impact of 5’UTR length, separating transcripts with long (>1,000 nucleotides) or short (<1,000 nucleotides) 5’UTRs. Transcripts with short 5’UTRs showed increased translation when asRNAs overlapped with the 5’UTR and CDS (**Figure 7D**). Moreover, transcripts with long 5’UTRs showed increased translation when asRNAs overlapped the 5’UTR and CDS or only the 5’UTR (**Figure 7D**). In contrast, asRNA overlapping with only the CDS or the 3’UTR had no impact on ribosome occupancy of the cognate sense protein coding transcripts (**Figure 7D**). Altogether, these findings highlight the tendency of asRNAs to enhance ribosome occupancy when overlapping the 5’UTR and CDS, particularly in transcripts with longer 5’UTRs. This underscores the important regulatory role of asRNAs and highlights a potentially widespread mechanism to facilitate translation of their cognate protein coding transcripts.

**Figure 7:**
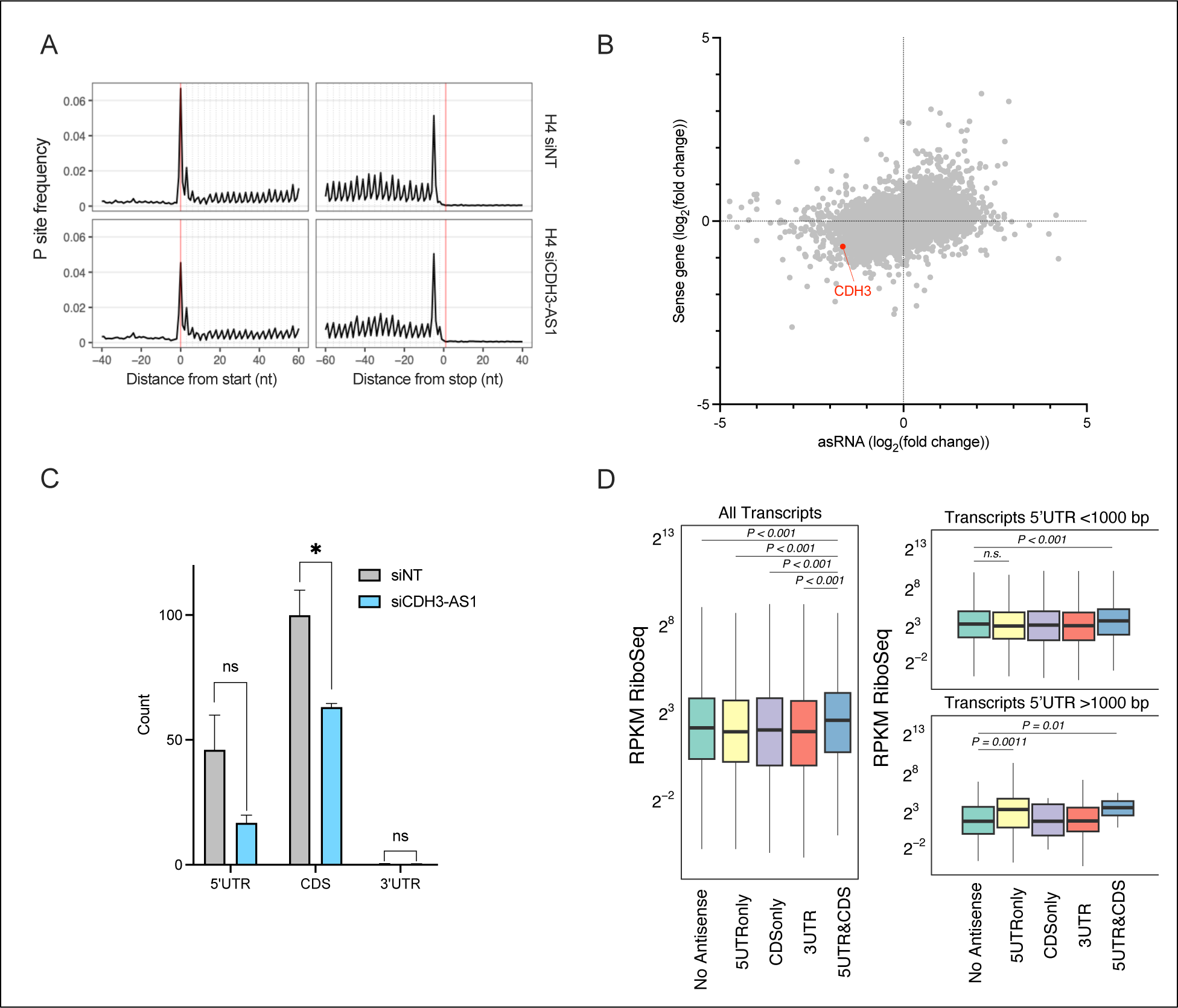
asRNA in the 5’UTR and CDS sequences influences ribosome occupancy and translation efficiency. (A) Metagene analysis of the density of 5’ read ends across all transcripts from annotated start codon to stop codon. (B) Integrated analysis of differential mRNA abundance (y-axis) and differential ribosome occupancy (x-axis). (C) Normalized read density in the 5’UTR, CDS, and 3’UTR of CDH3 in siNT and siCDH3-AS1 conditions. (D) Effects of antisense RNAs on average RPKM ribosome profiling read abundance of protein coding transcripts. * = p<0.05; ** = p<0.01; *** = p<0.001; ns = not significant.

## DISCUSSION

asRNAs are important regulators of gene expression, with potential impacts on cancer progression, including melanoma, positioning them as possible therapeutic targets. However, the mechanisms by which asRNAs exert their effects remain incompletely understood. In this study, we defined the profile of differentially expressed asRNA in melanoma and functionally characterized *CDH3-AS1*, which is significantly downregulated in melanoma. Our findings demonstrate that *CDH3-AS1* functions as a tumor suppressor by modulating the translational efficiency of P-cadherin. This work adds to our understanding of how asRNAs regulate their cognate sense genes and influence cancer development.

In this study, we identified strong downregulation of the *CDH3-AS1* asRNA and its cognate protein coding mRNA, *CDH3*, in melanoma. Our results indicate that the MAPK pathway, commonly activated in melanoma, leads to reduced expression of *CDH3-AS1* and *CDH3*. Both genes appear to be independently regulated by the MAPK pathway, as we observed that the expression of *CDH3-AS1* does not influence *CDH3* mRNA levels, and vice versa, suggesting the independent transcriptional regulation of these RNAs. Given the *CDH3-AS1* effect on the cognate sense *CDH3* mRNA, it is tempting to speculate that *CDH3* and *CDH3-AS1* are transcriptionally co-regulated to ensure efficient translation of P-cadherin. Moreover, a reduction in P-cadherin increases *CDH3- AS1* expression, indicating the existence of a transcriptional feedback mechanism to maintain P-cadherin translation. MAPK pathway hyperactivation, typically triggered by primary mutations in *BRAF*, *NRAS*, or *NF1* (42), is considered the initiating event in the malignant transformation of melanocytes. Thus, *CDH3-AS1* and *CDH3* are likely downregulated early during melanomagenesis, which is supported by our finding that BRAF^V600E^ expression reduces the levels of *CDH3-AS1* and *CDH3* in melanocytes, indicating tumor suppressive roles for both transcripts.

Previous studies have shown conflicting roles for P-cadherin, with both tumor suppressive (31–33) and oncogenic (34) functions described in melanoma. Our in vitro and in vivo findings support the tumor suppressive role of P-cadherin in melanoma. P-cadherin plays a role in calcium-dependent cell-cell adhesion (28) and forms heterologous cell junctions between melanocytes and keratinocytes (43). It is conceivable that a reduction of P-cadherin deregulates the crosstalk of transformed melanocytes with their microenvironment to facilitate melanomagenesis. However, keratinocytes may also influence P-cadherin levels in melanocytes through homotypic transcellular interaction of P-cadherin molecules in the skin microenvironment (43) and thus cell autonomous and non-autonomous mechanisms may regulate P-cadherin. Moreover, some studies suggested a negative correlation between *CDH3* expression and patient survival (35, 43); however, whether this is due to deregulated P-cadherin expression in melanoma cells remains to be elucidated and functionally evaluated in autochthonous mouse models. Overall, our results support that *CDH3* and *CDH3-AS1* possess tumor suppressive functions in melanoma: *CDH3* because it encodes P-cadherin to maintain crosstalk with the melanocyte microenvironment and *CDH3-AS1* because it facilitates the translation of P-cadherin.

Our study elucidates how *CDH3-AS1* increases P-cadherin protein levels. The most common mechanism of action reported for asRNAs is the regulation of its gene locus in cis (12, 44). This includes both the transcriptional regulation of the cognate sense gene and post-transcriptional influences on the mRNA encoded by the cognate sense gene. Using lentiviral and CRISPRa-mediated overexpression approaches, we found that *CDH3-AS1* influences P-cadherin expression more effectively when transcribed from its endogenous locus (CRISPRa approach), suggesting a function in cis. We show that *CDH3-AS1* does not affect the transcription of *CDH3* but rather *CDH3-AS1* interacts with *CDH3* mRNA. Moreover, lentiviral *CDH3-AS1* overexpression also moderately increases P-cadherin levels, suggesting that while the regulation of *CDH3* by *CDH3-AS1* is not strictly in cis, proximity of *CDH3-AS1* to the site of *CDH3* transcription increases the chance of interaction between the two transcripts. This is reminiscent of other lncRNAs acting in cis, and *CDH3-AS1* may therefore be described as a not “true” cis-acting lncRNA (45). Interestingly, both *CDH3-AS1* and *CDH3* are predominantly localized in the cytoplasm and silencing of *CDH3-AS1* with siRNA, which occurs in the cytoplasm, reduced P-cadherin translation. Taken together, these findings propose a model where *CDH3-AS1* interacts with *CDH3* in the nucleus near the site of transcription, followed by translocation of the asRNA/mRNA pair to the cytoplasm where *CDH3-AS1* stimulates the translation of P-cadherin through its interaction with *CDH3*.

We found that *CDH3-AS1* has no impact on *CDH3* intron 1 splicing, *CDH3* localization or turnover, or P-cadherin protein stability. Instead, we discovered putative secondary structures in the 5’UTR and early CDS of *CDH3* that robustly impair translation efficiency. Multiple studies have highlighted the importance of the 5’UTR of protein coding genes to regulate translational efficiency, in part due to the presence of secondary structures (46, 47). Interestingly, the RNA sequence in *CDH3* that contains the secondary structures is the region complementary to *CDH3- AS1* where the interaction of the two transcripts likely occurs. While our study demonstrates that *CDH3-AS1* robustly enhances P-cadherin levels, the exact mechanism of how *CDH3-AS1* facilitates the translation of *CDH3* requires further investigation. *CDH3-AS1* could affect *CDH3* translation in several possible ways. *CDH3-AS1* may promote the assembly of ribosomes on the 5’UTR, thereby enhancing translation. This could explain why the silencing of *CDH3-AS1* resulted in reduced ribosome occupancy on both the 5’UTR and CDS of *CDH3*. This process could either involve an upstream ORF (uORF) that directs translation (48) or be akin to “standby” ribosomes in prokaryotes that assemble upstream of a secondary structure and upon the temporary resolution of the secondary structure immediately begin translation (49). We showed that the 5’ region of *CDH3* harbors secondary structures that limit translation; however, asRNAs in prokaryotes may inhibit rather than stimulate translation by stand-by ribosomes (49).

Nevertheless, *CDH3-AS1* could promote the resolution of the *CDH3* secondary structures and thereby promote translation. This resolution could be mediated through the direct interaction of *CDH3-AS1* with *CDH3* or by the recruitment of RNA-binding proteins that aid in unwinding RNA. Indeed, *CDH3* translation is inhibited by Silvestrol and this effect is not rescued by *CDH3-AS1* overexpression, suggesting that *CDH3-AS1* requires helicase-mediated unwinding of *CDH3* to promote translation. Moreover, *CDH3-AS1* could act as a scaffold that recruits RNA-binding proteins (RBPs) to the 5’ region of *CDH3*. These RBPs could stimulate translation in multiple ways, for instance by promoting ribosome assembly, initiation complex formation, or looping of the poly A tail.

We made the noteworthy discovery that transcripts harboring asRNAs that overlap at the 5’ region exhibit increased translational efficiency. Interestingly, asRNAs that overlap in the CDS or 3’UTR but not the 5’UTR have no effect on translation dynamics, indicating that interaction of an asRNA with the 5’ region of a protein coding transcript is critical for this effect. Moreover, while transcripts with short 5’UTRs require overlap of the asRNAs with both the 5’UTR and parts of the CDS, asRNA overlap with only the 5’UTR of transcripts with longer 5’UTRs (>1,000 nucleotides) is sufficient to increase translation dynamics. This observation suggests that the length of overlapping sequence between asRNA and sense transcript determines the effects of translation, with longer complementary sequences likely having a stronger impact on translation. More work is needed to address questions regarding the regulatory impact of asRNA on the translation of their complementary protein coding transcripts. For instance, is the mechanism controlling *CDH3* translation shared with other asRNA/mRNA pairs? More specifically, if *CDH3-AS1* anneals with the 5’ region of *CDH3*, how would ribosomes assemble on double-stranded RNA? And if *CDH3- AS1* and *CDH3* already anneal in the nucleus, how would the temporary resolution of 5’UTR secondary structures be regulated in the cytoplasm? Nevertheless, our work highlights the functional importance of asRNA in regulating their cognate sense transcripts. Taken together, this study defines the differentially expressed asRNA profile in melanoma, characterizes the regulation of P-cadherin translation by *CDH3-AS1*, and proposes an asRNA-mediated mechanism of translational regulation of transcripts harboring overlapping asRNAs in their 5’UTRs.

## AUTHORS CONTRIBUTIONS

**M.C.**: Conceptualization, formal analysis, investigation, visualization, methodology, writing– original draft, writing–review and editing. **C.G.**: Formal analysis, investigation. **X.X.**: Investigation, writing–review and editing. **E.B.**: Formal analysis and investigation. **O.V.**: Investigation, writing– review and editing. **N.M.**: Investigation, **K.W.**: Investigation, **A.M.J.**: Formal analysis, investigation, writing–review and editing. **F.A.K.**: Conceptualization, formal analysis, supervision, funding acquisition, investigation, visualization, methodology, writing–original draft, project administration, writing–review and editing.

## ACKNOWLEDGMENTS

We are grateful to Karreth lab members for helpful discussions. We acknowledge the use of OpenAI’s ChatGPT to improve readability of the manuscript while all intellectual responsibility for the content remains with the authors.

