## Supplemental Figures for "CDH3-AS1 antisense RNA enhances P-cadherin translation and acts as a tumor suppressor in melanoma"

A

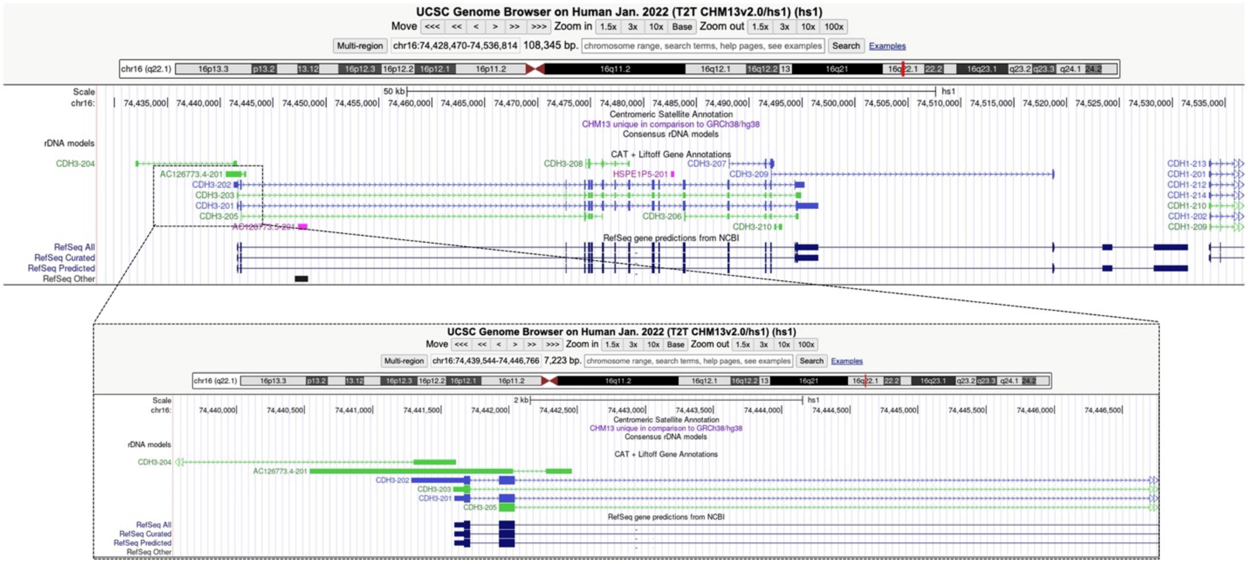

B

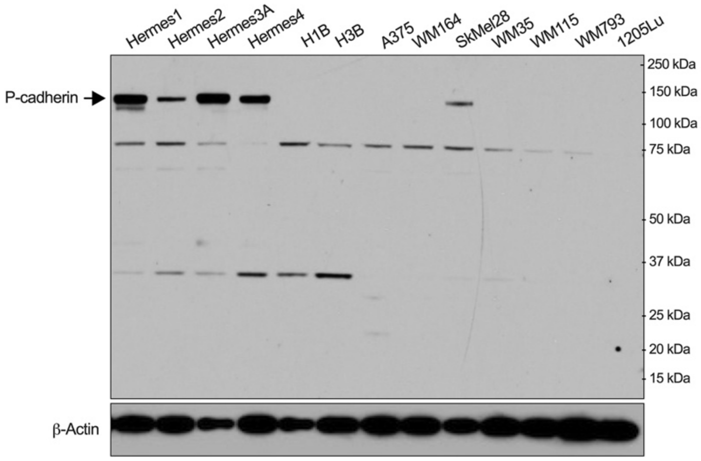


**Supplementary Figure 1: Genomic organization of the CDH3 locus and P-Cadherin size.**

(A) UCSC Genome Browser view of the *CDH3* genomic locus, highlighting the overlap between *CDH3-AS1* and *CDH3*. (B) Western blot analysis showing P-cadherin protein levels (120 kDa) alongside additional bands at 75 kDa and 30 kDa.

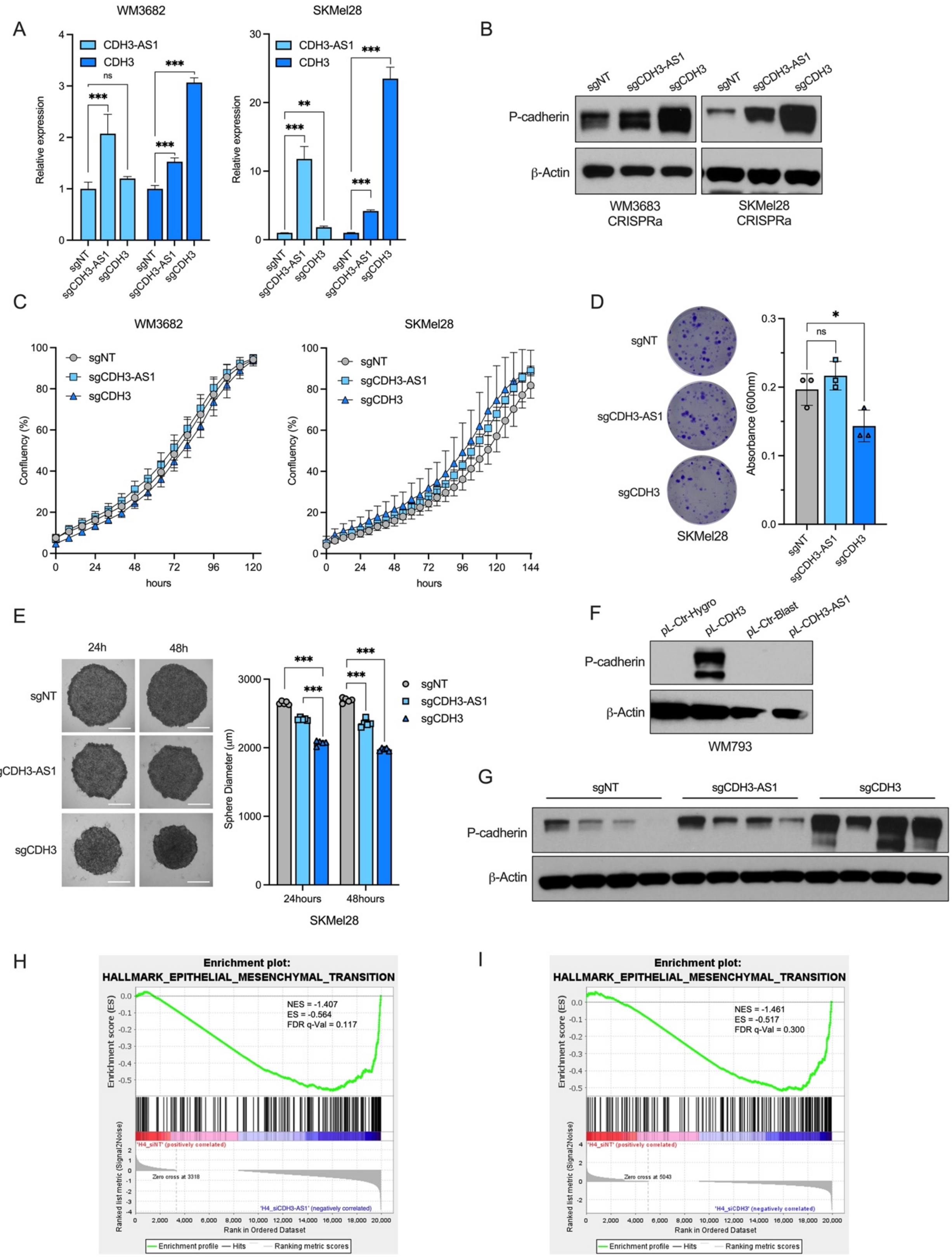
A
WM3682
SKMel28
CDH3-AS1
CDH3
Relative expression
ns
***
**
sgNT
sgCDH3-AS1
sgCDH3
B
P-cadherin
β-Actin
WM3683
CRISPRa
SKMel28
CRISPRa
C
WM3682
SKMel28
Confluency (%)
hours
D
SKMel28
Absorbance (600nm)
*
E
24h
48h
Sphere Diameter (μm)
24hours
48hours
SKMel28
F
pL-Ctr-Hygro
pL-CDH3
pL-Ctr-Blast
pL-CDH3-AS1
WM793
G
H
Enrichment plot:
HALLMARK_EPITHELIAL_MESENCHYMAL_TRANSITION
NES = -1.407
ES = -0.564
FDR q-Val = 0.117
Enrichment score (ES)
'H4_siNT' (positively correlated)
Zero cross at 3318
'H4_siCDH3-AS1' (negatively correlated)
Ranked list metric (Signal2Noise)
Rank in Ordered Dataset
Enrichment profile
Hits
Ranking metric scores
I
Enrichment plot:
HALLMARK_EPITHELIAL_MESENCHYMAL_TRANSITION
NES = -1.461
ES = -0.517
FDR q-Val = 0.300
'H4_siNT' (positively correlated)
Zero cross at 5043
'H4_siCDH3' (negatively correlated)

**Supplementary Figure 2: *CDH3-AS1* and *CDH3* impact slow aggregation and EMT pathway.** (A) RT-qPCR analysis showing overexpression of *CDH3-AS1* and *CDH3* in WM3682 and SKMel28 cells using CRISPRa. (B) Western blot analysis of P-cadherin levels in WM3682 and SKMel28 cells with CRISPRa. (C) Proliferation assay measuring relative confluence of WM3682 and SKMel28 cells overexpressing *CDH3-AS1* or *CDH3*. (D) Low-density colony formation assay (representative images on left, quantification on right) in SKMel28 cells overexpressing *CDH3-AS1* or *CDH3*. (E) Slow aggregation assay (representative images on left, quantification on right) in SKMel28 cells with stable overexpression of *CDH3-AS1* or *CDH3*. (F) Western blot analysis of P-cadherin levels in WM793 cells with stable transfection of pL-Ctr (Hygro or Blast), pL-CDH3 or pL-CDH3-AS1. (G) Western blot analysis of P-cadherin levels in tumors collected from NSG mice xenografted with WM3682 cells overexpressing *CDH3-AS1* or *CDH3*. (H-I) Enrichment plots of the epithelial-mesenchymal transition hallmark pathway in Hermes4B cells upon silencing of *CDH3-AS1* (H) or *CDH3* (I). * = $p < 0.05$; ** = $p < 0.01$; *** = $p < 0.001$; ns = not significant.

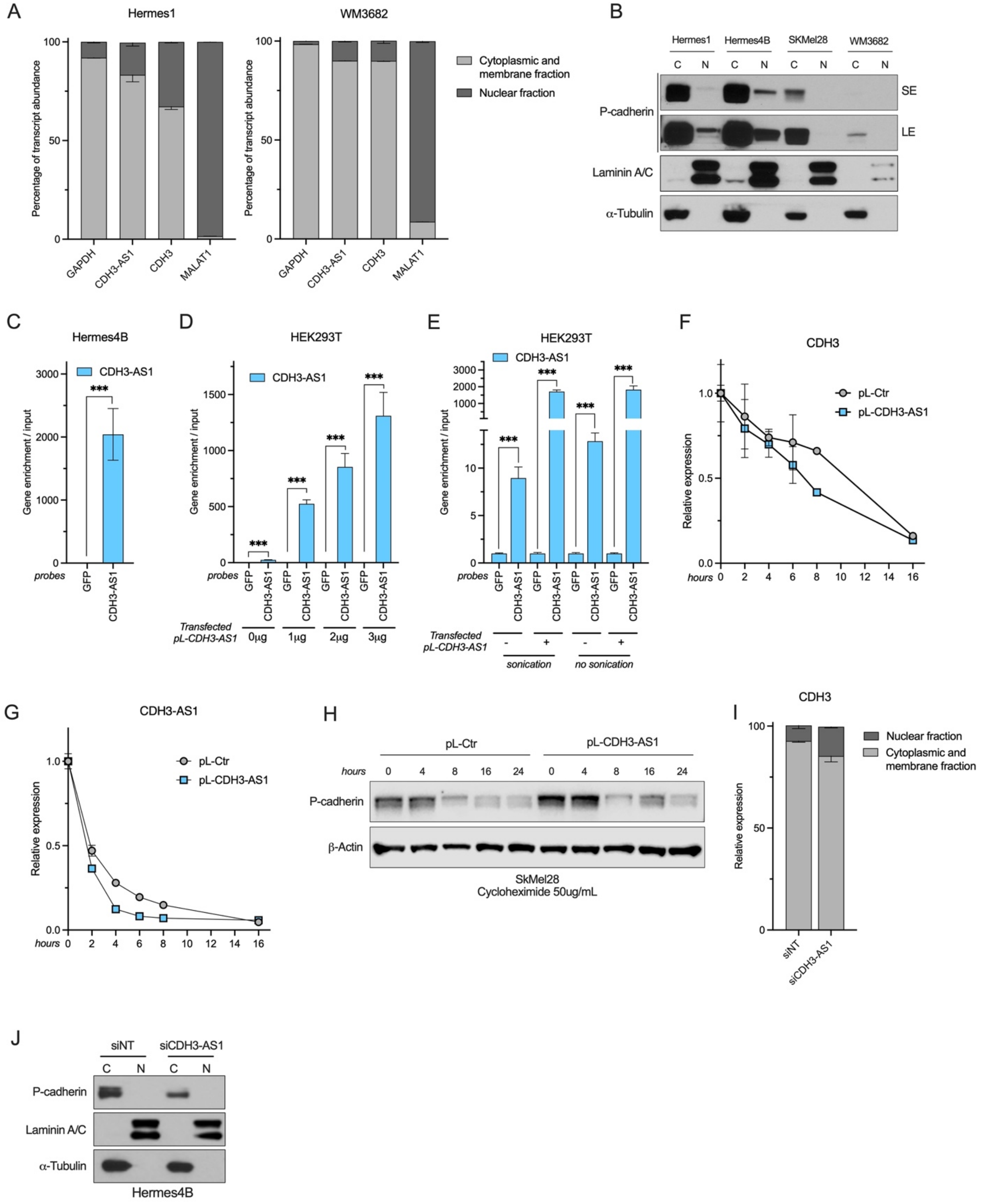
A
Hermes1
WM3682
Percentage of transcript abundance
100
50
0
GAPDH
CDH3-AS1
CDH3
MALAT1
Cytoplasmic and membrane fraction
Nuclear fraction
B
Hermes1
Hermes4B
SKMel28
WM3682
C
N
P-cadherin
SE
LE
Laminin A/C
α-Tubulin
C
Hermes4B
CDH3-AS1
Gene enrichment / input
3000
2000
1000
0
probes
GFP
***
D
HEK293T
1500
1000
500
Transfected pL-CDH3-AS1
0μg
1μg
2μg
3μg
E
HEK293T
3000
2000
1000
10
5
sonication
no sonication
-
+
F
CDH3
pL-Ctr
pL-CDH3-AS1
Relative expression
1.0
0.5
0.0
hours
0
2
4
6
8
10
12
14
16
G
CDH3-AS1
H
pL-Ctr
pL-CDH3-AS1
hours
0
4
8
16
24
P-cadherin
β-Actin
SkMel28
Cycloheximide 50ug/mL
I
CDH3
Nuclear fraction
Cytoplasmic and membrane fraction
Relative expression
siNT
siCDH3-AS1
J
siNT
siCDH3-AS1
P-cadherin
Laminin A/C
α-Tubulin
Hermes4B

**Supplementary Figure 3: Subcellular localization and functional analysis of *CDH3-AS1* in regulating *CDH3* expression and P-Cadherin translation.**

(A-B) Nuclear and cytoplasmic fractionation in Hermes4B cells followed by RT-qPCR to detect *GAPDH* (cytoplasmic control), *CDH3-AS1*, *CDH3*, and *NEAT1* (nuclear control) transcripts (A) or Western blot to detect P-cadherin, Laminin A/C (nuclear control), and α-Tubulin (cytoplasmic control) protein levels (B). (C-E) Enrichment of *CDH3-AS1* RNA in RNA pull-down experiments using biotinylated *CDH3-AS1* probes in Hermes4B cells (C), HEK293T cells co-transfected with equal amounts of pL-CDH3 plasmid and increasing amounts of pL-CDH3-AS1 plasmid (D) and with or without sonication (E). (F-G) Relative expression of *CDH3* (F) or *CDH3-AS1* (G) at different time points after actinomycin D treatment in cells overexpressing *CDH3-AS1* or control cells. (H) Western blot analysis of P-cadherin protein levels at different time points following cycloheximide treatment in cells overexpressing *CDH3-AS1* or control cells. (I-J) Nuclear and cytoplasmic fractionation of Hermes4B cells upon *CDH3-AS1* knockdown, followed by RT-qPCR (I) and Western blot (J) to detect *CDH3* mRNA and P-cadherin protein levels, respectively. SE: Short exposure; LE: Long exposure. *** = $p < 0.001$.

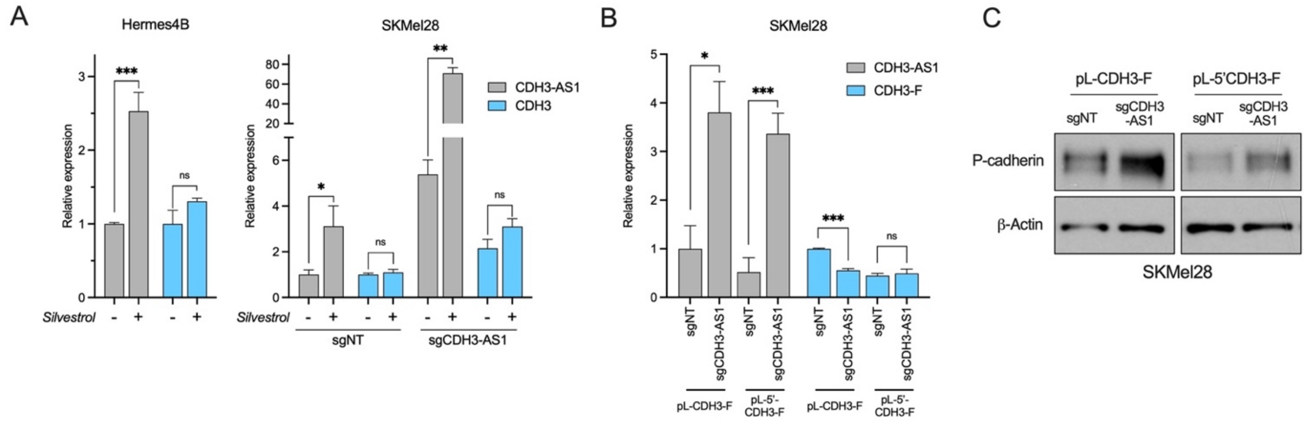


**Supplementary Figure 4: *CDH3-AS1* feedback loop under Silvestrol treatment.**

(A) RT-qPCR analysis showing the relative expression levels of *CDH3-AS1* and *CDH3* in Hermes4B and SKMel28 cells overexpressing *CDH3-AS1* with and without Silvestrol treatment. (B) RT-qPCR analysis showing the expression of *CDH3-AS1* and *CDH3-F* (CDH3-FLAG) in SKMel28 cells with stable overexpression of *CDH3-AS1* and transient transfection of pL-CDH3-F or pL-5'-CDH3-F constructs. (C) Western blot analysis of P-cadherin protein levels in SKMel28 cells with stable *CDH3-AS1* overexpression and transient transfection of pL-CDH3-F or pL-5'-CDH3-F constructs. * = $p < 0.05$; ** = $p < 0.01$; *** = $p < 0.001$; ns = not significant.

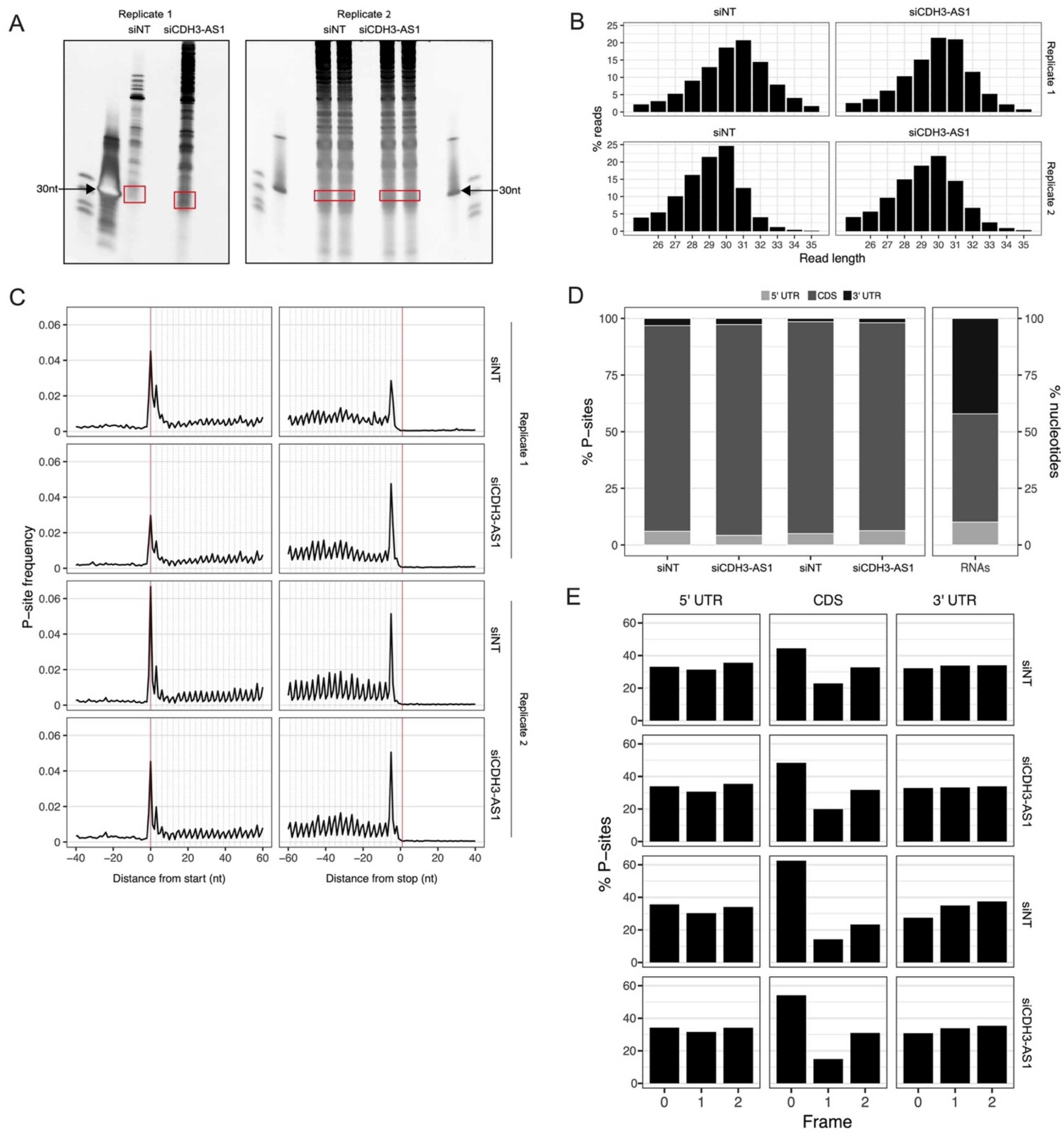


**Supplementary Figure 5: Ribosome profiling.**

(A) Size selection of ribosome protected fragments of RNA (RPFs) with SDS-PAGE. The excised area used for library preparation is highlighted in red. (B) Length distributions of sequenced RNA. (C) Metagene analysis of tri-nucleotide periodicity within the CDS for all samples. (D) Percentage of P-sites that mapped to the 5'UTR, CDS, and 3'UTR for each sample. To the right, the percentage of nucleotides which map to the 5'UTR, CDS, and 3'UTR across all RNA. (E) Percentage of P-sites that align to reading frame 0, +1, or +2 for the 5'UTR, CDS, and 3'UTR.
